# SIRPα controls CD47-dependent platelet clearance in mice and humans

**DOI:** 10.1101/2023.12.09.570874

**Authors:** Maia Shoham, Ying Ying Yiu, Paige S. Hansen, Aanya Subramaniam, Martin Broberg, Eric Gars, Tal Raveh, FinnGen, Irving L. Weissman, Nasa Sinnott-Armstrong, Anandi Krishnan, Hanna M. Ollila, Michal Caspi Tal

## Abstract

Over the last decade, more data has revealed that increased surface expression of the “don’t eat me” CD47 protein on cancer cells plays a role in immune evasion and tumor progression, with CD47 blockade emerging as a new therapy in immuno-oncology. CD47 is critical in regulating cell homeostasis and clearance, as binding of CD47 to the inhibitory receptor SIRPα can prevent phagocytosis and macrophage-mediated cell clearance. The purpose of this study was to examine the role of the CD47-SIRPα signal in platelet homeostasis and clearance. Therapeutic reagents targeting the CD47-SIRPα axis are very promising for treatment of hematologic malignancies and solid tumors, but lead to transient anemia or thrombocytopenia in a subset of patients. We found that platelet homeostatic clearance is regulated through the CD47-SIRPα axis and that therapeutic blockade to disrupt this interaction in mice and in humans has a significant impact on platelet levels. Furthermore, we identified genetic variations at the *SIRPA* locus that impact platelet levels in humans such that higher *SIRPA* gene expression is associated with higher platelet levels. *SIRPA* expression at either end of the normal range may affect clinical outcomes of treatment with anti-CD47 therapy.

**Key points:**

- Platelet homeostasis is regulated through the CD47-SIRPα axis and therapeutic blockade to disrupt this interaction impacts platelet levels
- Common genetic variants at *SIRPA* locus associate with platelet levels

## Introduction

CD47, also known as integrin-associated protein (IAP), is a transmembrane glycoprotein expressed on hematopoietic and non-hematopoietic cells that, when bound to the highly polymorphic immune inhibitory receptor Signal Regulatory Protein Alpha (SIRPα), reduces phagocytosis and macrophage-mediated clearance of cells. Effectively, macrophages expressing SIRPα receive a ‘don’t eat me’ signal upon binding to CD47, as reviewed by Barclay and Ven den Berg (2014) ^1^. The potency of the ‘don’t eat me’ signal depends on several factors, including the relative level of surface CD47 ^2^. In the case of pre-cancerous cells, the up-regulation of CD47 contributes to immune evasion of potentially harmful cells. Cancer cells upregulate CD47 expression to evade immune clearance and the level of expression may correlate with poor clinical outcomes in some cancers ^3^. In contrast, CD47 expression declines with cell age, notably serving as a biological timer for red blood cell (RBC) clearance, whereby loss of CD47 expression on mature red blood cells marks aged erythrocytes heading for clearance ^4^. Because of its role in regulating cell clearance, CD47 has been targeted in immunotherapeutic treatment approaches for hematologic malignancies and solid tumors ^5–7^. Inhibition of CD47’s anti-phagocytic signaling is a promising avenue for cancer immunotherapy ^5^. The CD47-SIRPα axis has also been implicated in the regulation of homeostatic levels of different immune cell populations. Previous studies demonstrated that CD47 blockade *in vivo* resulted in enhanced erythrophagocytosis leading to transient anemia ^6^. Furthermore, in a mouse model of induced colitis, mice lacking SIRPα or CD47 also developed anemia, followed by compensatory extramedullary erythropoiesis leading to splenomegaly ^8^. Interestingly, platelet levels have also been shown to be regulated by the CD47-SIRPα axis ^9^. With several human clinical trials evaluating the blockade of the CD47-SIRPα axis completed or in progress, there is considerable focus on preventing treatment-induced anemia (**Table 1**). In many of these trials, transient thrombocytopenia has also been noted in some patients. However, it is not known why some individuals develop anemia or thrombocytopenia, while others do not.

**Table 1.**
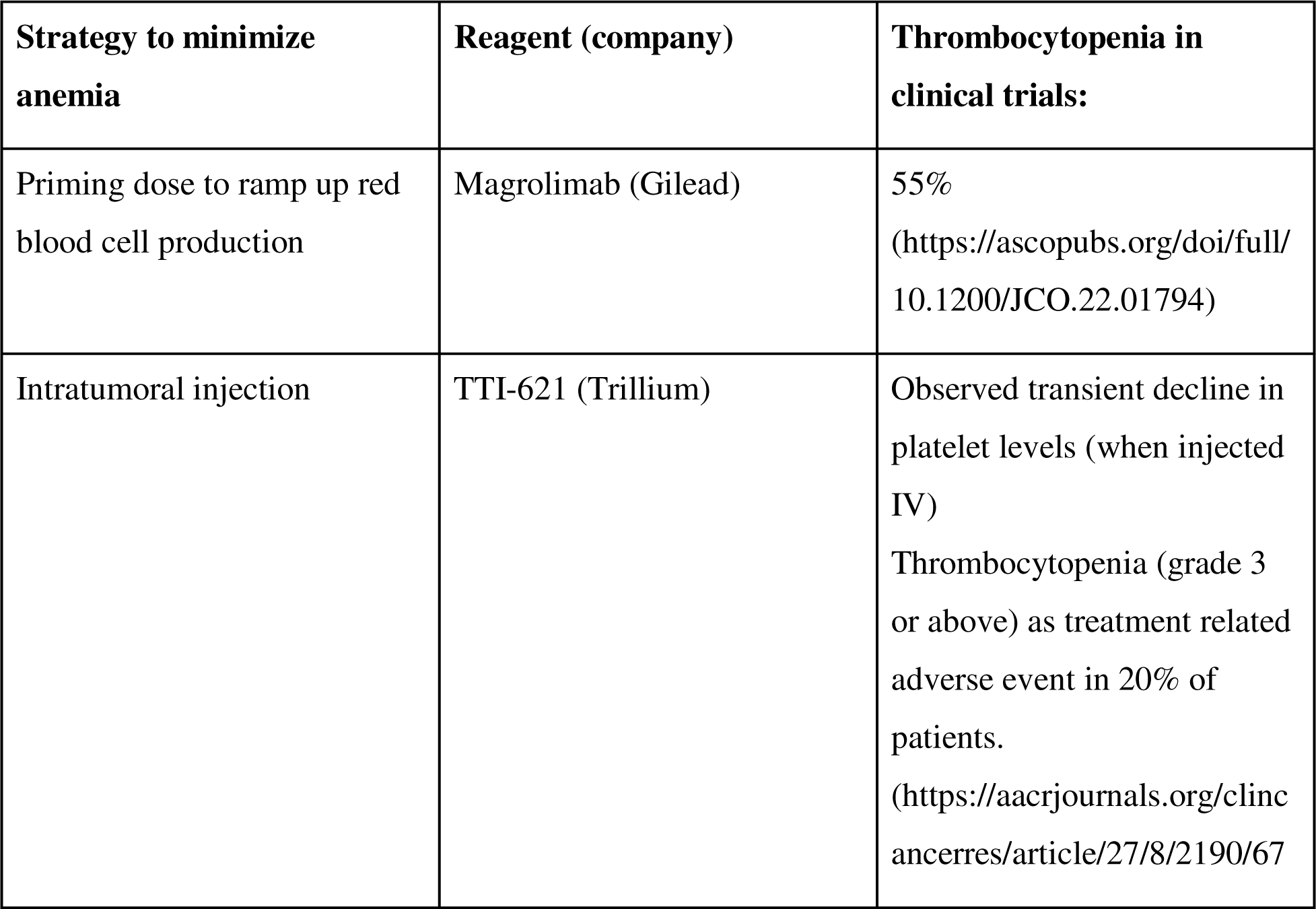
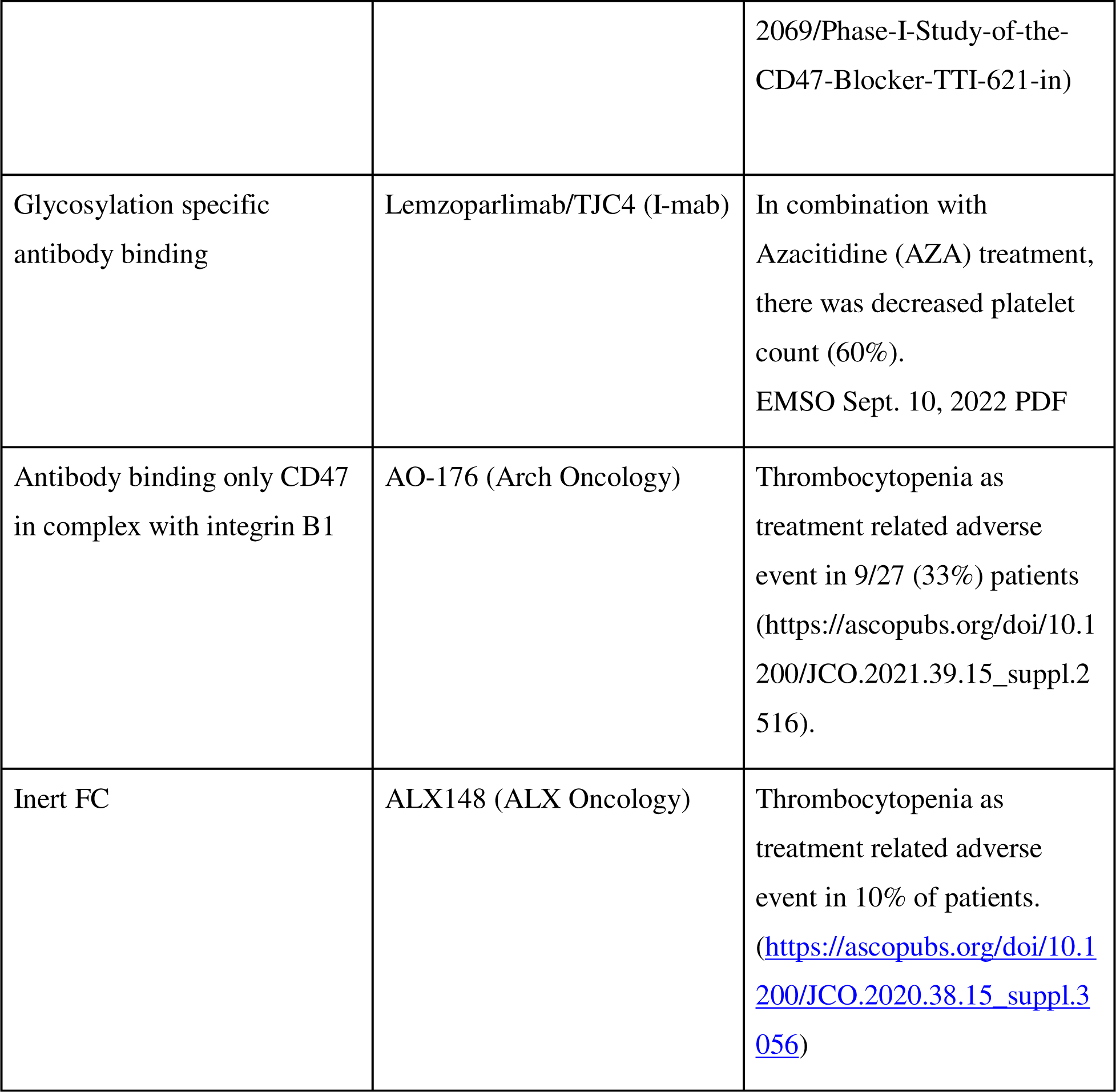
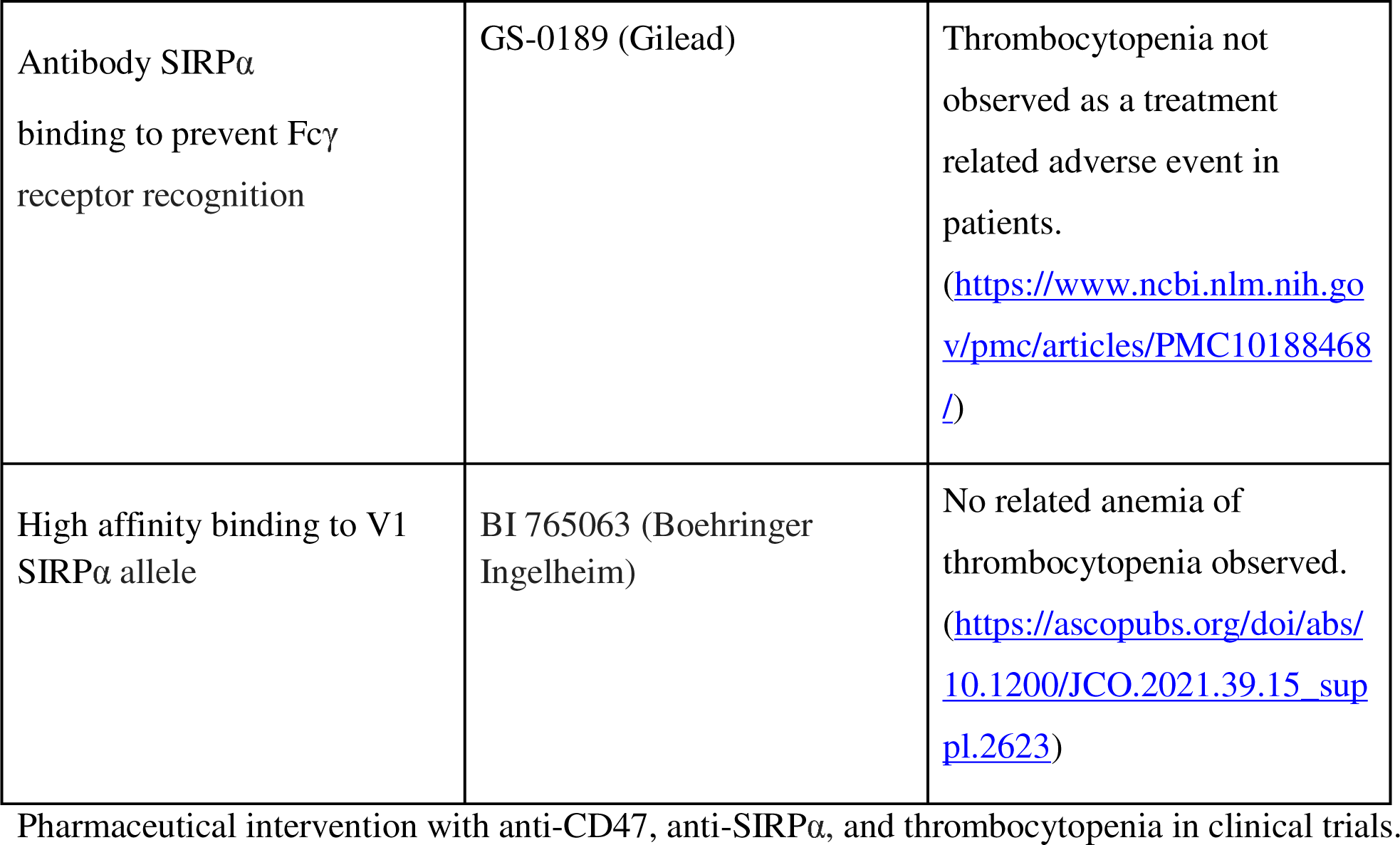

Previously, genetic association studies have provided insight into the pathophysiology of a variety of immune cell traits ^10^. These earlier studies have identified specific genetic variants that affect platelet and erythrocyte traits in humans. Similarly, variant-to-expression analyses have provided insight into how genetic variants contribute to changes in RNA levels^11^ and protein expression ^12^, with possible implications for disease mechanisms. Because of the known connection between CD47/SIRPα and RBC homeostasis and this axis’ association with platelets ^5,9^, we sought to understand the effect of *in vivo* CD47 blockade on platelet clearance and the broader association between *SIRPA* polymorphisms and platelet phenotypes.

## Materials and Methods

For clarity of nomenclature, we use *SIRPA* for the gene name and SIRPα for the protein name throughout the text.

### *In vivo* CD47 Blockade

All experiments were carried out in accordance with ethical care guidelines set by the Stanford University Administrative Panel on Laboratory Animal Care (APLAC). Female mice were used for all studies. Investigators were not blinded for animal studies.

C57BL/6 mice were injected intraperitoneally with a priming dose of 100 µg, followed by 500µg treatment doses of anti-CD47 MIAP410 (Bio X Cell) antibody, IgG isotype control or PBS every 48 hours for 2 weeks. After euthanasia, cardiac puncture was performed and spleens were collected. Blood was collected in diagnostic tubes containing EDTA and submitted to the veterinary diagnostic laboratory at Stanford for CBC. Spleens were preserved in 4% paraformaldehyde for 12 hours at 4 degrees overnight, then transferred to 70% ethanol for submission to HistoWiz. They were cut at 5 µm thickness and stained for hematoxylin and eosin (H&E) and Ter119.

### Meta-analysis of earlier GWAS

We performed a meta-analysis in three cohorts with total N=709,883. These cohorts include meta-analysis of previously published summary statistics from the UK Biobank (N=444,382, accessed through https://alkesgroup.broadinstitute.org/LDSCORE/all_sumstats/), Biobank Japan (N=108,208) ^13^ and Blood Cell Consortium (N=157,293) ^14^. We used fixed effects meta-analysis as implemented in META v1.7 ^15^.

### FinnGen

FinnGen is a large-scale research study that aims to genotype 500,000 Finnish participants recruited from hospitals as well as prospective and retrospective epidemiological and disease-based cohorts. These data are combined with longitudinal registries that record phenotypes and health events (including ICD-based diagnosis) over the entire lifespan including the National Hospital Discharge Registry (inpatient and outpatient), Causes of Death Registry, the National Infectious Diseases Registry, Cancer Registry, Primary Health Care Registry (outpatient) and Medication Reimbursement Registry.

This study used data from FinnGen Data Freeze 9, which comprises 377,277 individuals. For the purposes of our research, we used registered sleep medication purchases (which constitute a prescription-only medication in Finland) dating from 1995. The primary indication for sleep medication prescription in Finland is sleep problems, and of these, non-benzodiazepine drugs are used for insomnia treatment ^16^.

### FinnGen ethics statement

Patients and control subjects in FinnGen provided informed consent for biobank research, based on the Finnish Biobank Act. Alternatively, separate research cohorts, collected before the Finnish Biobank Act came into effect (in September 2013) and before start of FinnGen (August 2017), were collected based on study-specific consents and later transferred to the Finnish biobanks after approval by Fimea (Finnish Medicines Agency), the National Supervisory Authority for Welfare and Health. Recruitment protocols followed the biobank protocols approved by Fimea. The Coordinating Ethics Committee of the Hospital District of Helsinki and Uusimaa (HUS) statement number for the FinnGen study is Nr HUS/990/2017.

The FinnGen study is approved by the Finnish Institute for Health and Welfare (permit numbers: THL/2031/6.02.00/2017, THL/1101/5.05.00/2017, THL/341/6.02.00/2018, THL/2222/6.02.00/2018, THL/283/6.02.00/2019, THL/1721/5.05.00/2019 and THL/1524/5.05.00/2020), Digital and population data service agency (permit numbers: VRK43431/2017-3, VRK/6909/2018-3, VRK/4415/2019-3), the Social Insurance Institution (permit numbers: KELA 58/522/2017, KELA 131/522/2018, KELA 70/522/2019, KELA 98/522/2019, KELA 134/522/2019, KELA 138/522/2019, KELA 2/522/2020, KELA 16/522/2020), Findata permit numbers THL/2364/14.02/2020, THL/4055/14.06.00/2020,,THL/3433/14.06.00/2020, THL/4432/14.06/2020, THL/5189/14.06/2020, THL/5894/14.06.00/2020, THL/6619/14.06.00/2020, THL/209/14.06.00/2021, THL/688/14.06.00/2021, THL/1284/14.06.00/2021, THL/1965/14.06.00/2021, THL/5546/14.02.00/2020 and Statistics Finland (permit numbers: TK-53-1041-17 and TK/143/07.03.00/2020 (earlier TK-53-90-20)).

The Biobank Access Decisions for FinnGen samples and data utilized in FinnGen Data Freeze 7 include: THL Biobank BB2017_55, BB2017_111, BB2018_19, BB_2018_34, BB_2018_67, BB2018_71, BB2019_7, BB2019_8, BB2019_26, BB2020_1, Finnish Red Cross Blood Service Biobank 7.12.2017, Helsinki Biobank HUS/359/2017, Auria Biobank AB17-5154 and amendment #1 (August 17 2020), Biobank Borealis of Northern Finland_2017_1013, Biobank of Eastern Finland 1186/2018 and amendment 22 § /2020, Finnish Clinical Biobank Tampere MH0004 and amendments (21.02.2020 & 06.10.2020), Central Finland Biobank 1-2017, and Terveystalo Biobank STB 2018001.

### PheWAS

PheWAS analysis using FinnGen release 9 data for individual endpoints as predefined by the FinnGen project for coagulation and ischemic stroke. We also used Open Targets Genetics platform (https://genetics.opentargets.org/) ^17^ to explore association of *SIRPA* risk variants with other traits and biomarker GWAS.

### Mendelian Randomization

The meta-platelets summary statistics for effect size, standard error and P-values were used to find significant lead variants (P-value <5×10^−8^) using FUMA platform ^18^. Lead variants were subsequently used in a two-sample bidirectional Mendelian Randomization (MR) analysis. We estimated the effect from platelets on stroke and coronary artery disease using previous GWAS as identified from the GWAS catalog (June 2020 version, see Supplementary information for specific catalog IDs) ^19^. The MR was performed with the twosamplemr R package ^20^. Significant causality was explored based on Inverse Variance Weighting (IVW) statistical method (P-value < 0.05), and effect estimate was compared with estimates from MR Egger, IVW, Weighted Median, Simple Mode and Weighted Mode methods. Pleiotropy was assessed using MR Egger intercept.

### Human Subjects

For platelet RNA studies, study approval was provided by the Stanford University Institutional Review Board (#18329). We collected blood from patients with myeloproliferative neoplasms, MPNs (specifically essential thrombocythemia, ET in this study) enrolled in the Stanford University and Stanford Cancer Institute Hematology Tissue Bank between December 2017-2020. All MPN ET patient peripheral blood samples were obtained under written informed consent and were fully anonymized. All relevant ethical regulations were followed.

Eligibility criteria included age ≥18 years and Stanford MPN clinic diagnosis of essential thrombocythemia (defined using the consensus criteria at the time of this study). For healthy controls, blood was collected from eleven asymptomatic adult donors selected at random from the Stanford Blood Center. All donors were asked for consent for genetic research. For both ET patients and healthy controls, blood was collected into acid citrate-dextrose (ACD, 3.2%) sterile yellow-top tubes (Becton, Dickinson and Co.) and platelets were isolated by established ^21,22,23,24^ purification protocols. Blood was processed within 4 h of collection for all samples. The time from whole blood collection to platelet isolation was similar between healthy donors and MPN patients. For next generation RNA-sequencing (RNA-seq), 1×10^9^ isolated platelets lysed in Trizol were processed following established procedures ^21,22,25^ for RNA isolation (RNA integrity numbers >7.0), extraction and library preparation. Twelve pooled samples with individual indices were run on an Illumina HiSeq 4000 (Patterned flow cell with Hiseq4000 SBS v3 chemistry) as 2 X 75bp paired end sequencing with a coverage goal of 40M reads/sample. Resultant platelet transcriptomic data was library-size-corrected, variance-stabilized, and log2-transformed using the R package DESeq2 ^26^.

## Results

Thrombocytopenia emerges as a common adverse event across distinct CD47 blockade reagents, despite varying approaches aimed at minimizing anemia resulting from CD47-mediated red blood cell clearance (Table 1). Due to the role of the CD47-SIRPα axis in cell clearance and platelet homeostasis, we sought to investigate the possibility that blocking CD47 could directly trigger thrombocytopenia.

### CD47 blockade *in vivo* induces murine splenomegaly, splenic red pulp expansion from erythropoiesis, and increased megakaryocytes

Prior studies have shown that sustained CD47 blockade *in vivo* is most effective and survivable after an initial dose of a smaller quantity of the antibody is given, termed the priming dose. This priming dose triggers compensatory erythropoiesis and prevents fatal anemia. To investigate the effects of CD47 blockade in a murine model, we injected C57BL/6 mice with a priming dose of either PBS, IgG isotype control, or anti-mouse CD47 antibody, MIAP410, two days before commencing treatment. We then injected PBS, IgG, or MIAP410 every two days for two weeks. Upon dissection, splenomegaly was grossly apparent in specimens who received anti-CD47 mAb treatment (Fig. 1A). Histological examination showed expansion of red pulp due to extramedullary erythropoiesis (Fig. 1B). Furthermore, megakaryocyte infiltration into the spleen was observed (Fig. 1C), suggesting that the murine spleen becomes a site of increased platelet production in the setting of CD47 blockade.

**Figure 1.**
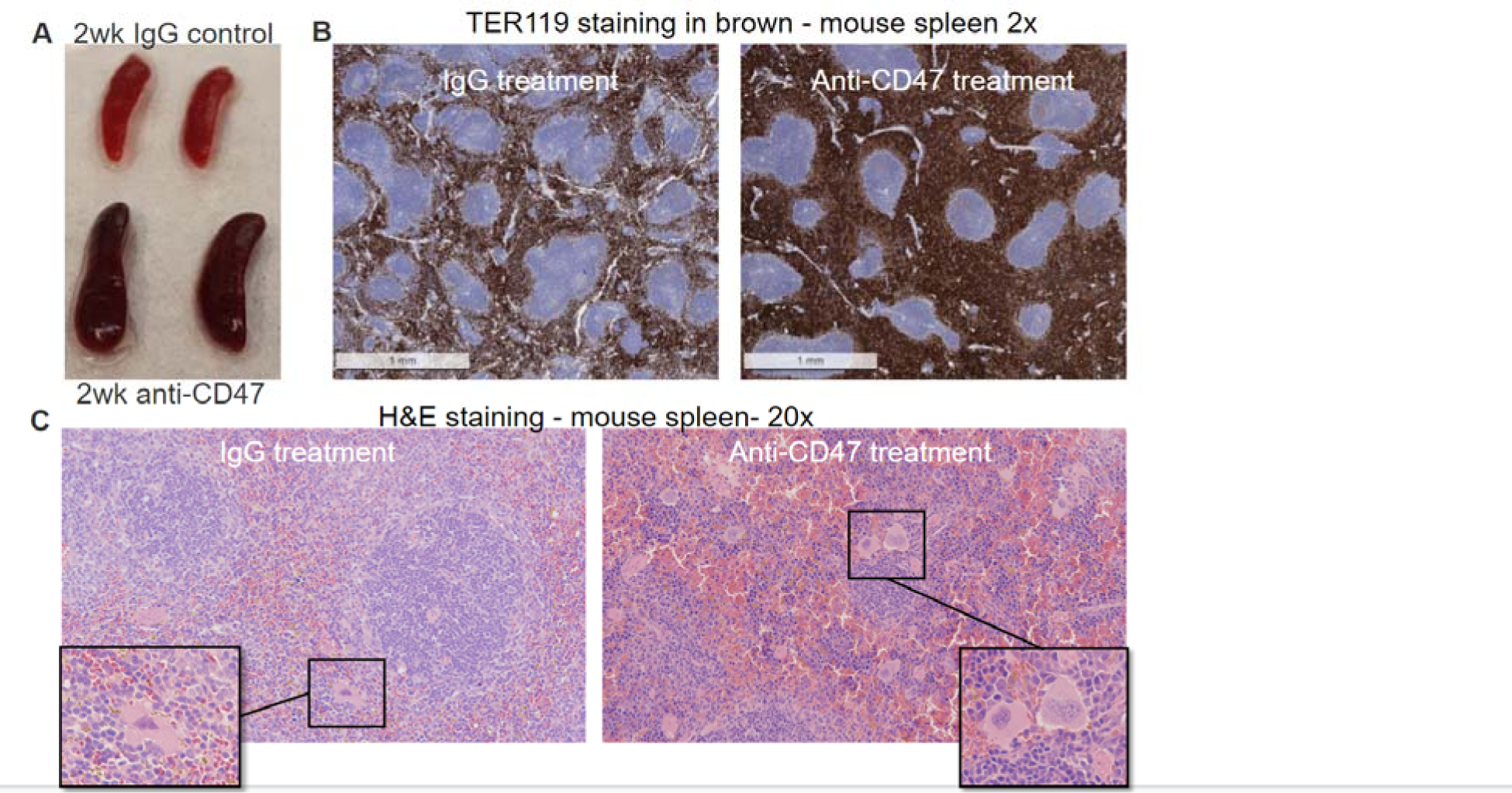
CD47 blockade induces splenomegaly, splenic red pulp expansion from erythropoiesis, and increased megakaryocytes. *In vivo,* CD47 blockade leads to splenomegaly with extramedullary erythropoiesis. Mice were treated with MIAP410 (clone-3 anti-CD47) for 2 weeks. (A) Spleens were removed and photographed with visible increase in size and pigment of spleens from animals who received the CD47 blockade treatment (bottom) compared to the IgG control treated animals (top). (B) Spleen sections 5uM thick were stained with anti-TER119 demonstrating expansion of erythroid cells in the red pulp of the spleen. (C) Hematoxylin and eosin (H&E) staining of interfollicular spaces of spleens demonstrates that the white pulp of th spleen is diminished in anti-CD47-treated mice, with the space now devoted to extramedullary erythropoiesis and platelet production, with significant increase in megakaryocytes evident in anti-CD47-treated mice (representative megakaryocytes shown in the boxes). (H&E) images at 200 µM resolution.

### Anti-CD47 treatment increases platelet clearance

Platelets use CD47 to evade phagocytosis by macrophages ^9^. To understand the effects of CD47 blockade on platelet homeostasis with greater precision, we collected mouse blood via cardiac puncture at the time of euthanasia and submitted those samples for complete blood count (CBC). Mice treated with MIAP410 had approximately three-fold fewer platelets than controls treated with either PBS or IgG isotype control (p = 0.0003, p = 0.0008, Fig. 2A). In addition to a reduction in the absolute number of platelets, the remaining platelets were approximately 12% larger in volume than controls (p = 0.0079, Fig. 2B MPV). There is an inverse relationship between MPV and platelet count (Fig. 2A and 2B), which has also been observed in human clinical data ^27^. P-LCR, which represents the percentage of platelets larger than 12fL, increased in treated mice (Fig. 2B P-LCR). Absolute reticulocyte count was also significantly increased in the treated mice (p = 0.0147, Fig. 2C), which demonstrates the hematopoietic system’s concerted efforts to compensate for the anemic and thrombocytopenic effects of anti-CD47 treatment. Based on maturation stage, reticulocytes can be classified into subtypes, where high fluorescence reticulocytes (HFR) and medium fluorescence reticulocytes (MFR) are immature and low fluorescence reticulocytes (LFR) are mature ^28^. In the treated mice, immature reticulocyte fraction (IRF) significantly increased (p = 0.0011, Fig. 2D), and LFR decreased (p = 0.0011, Fig. 2D), showing the effect of CD47 blockade with reticulocyte maturation.

**Figure 2.**
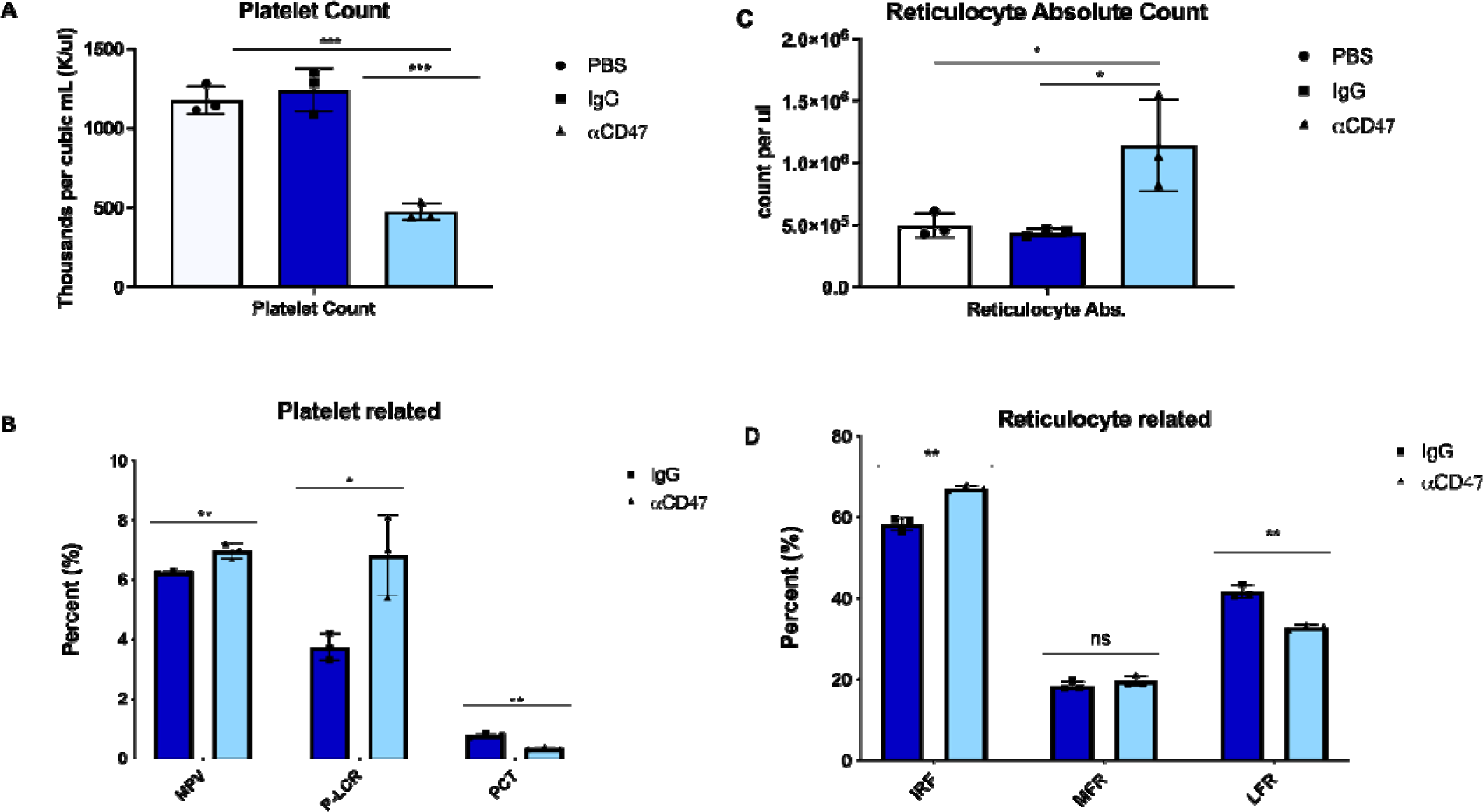
Platelets are reduced in number but increases in volume upon CD47 blockade in vivo. Complete blood counts were run after 2wks of anti-CD47 blockade in C57/B6 mice. A) Platelet count shown in thousands per cubic mL. B) Platelet related percentages shown a indicated, MPV: mean platelet volume (average platelet size), P-LCR: platelet large cell ratio, PCT: volume occupied by platelets in blood as percentage C) Reticulocyte absolute count shown per microliter. D) Reticulocyte related percentages shown as indicated, IRF: immature reticulocyte fraction, MFR: middle-fluorescence reticulocytes, LFR: low-fluorescence reticulocytes. CBC counts were run on two separate comparisons of CD47 blockade, one representative experiment is shown n=3 mice per group, significance of unpaired T-test comparisons shown as *(*p* ≤*.05, **p* ≤*.01, ***p* ≤*.005)*.

### Variation at *SIRPA* affects platelet levels in humans

Previous genome-wide association studies have implied that platelet levels are modulated by common genetic risk factors, including even SIRPA ^29^. We therefore performed a meta-analysis of N=752,910 individuals using publicly available GWAS evidence for 645 independent loci (Supplementary information). The significant loci included *BBX-CD47* and *SIRPA*. Specifically, we discovered GWAS associations at the 3’ end of *CD47,* close to the *BBX* gene (rs167924, beta = −0.018, p= 7.40e-23; Fig. 3A). In addition, we discovered that variants in the 3’UTR of *SIRPA* (rs3197744 beta = −0.023, p = 9.88e-32, Fig. 3B) are associated with the amount of platelets in those individuals.

**Figure 3.**
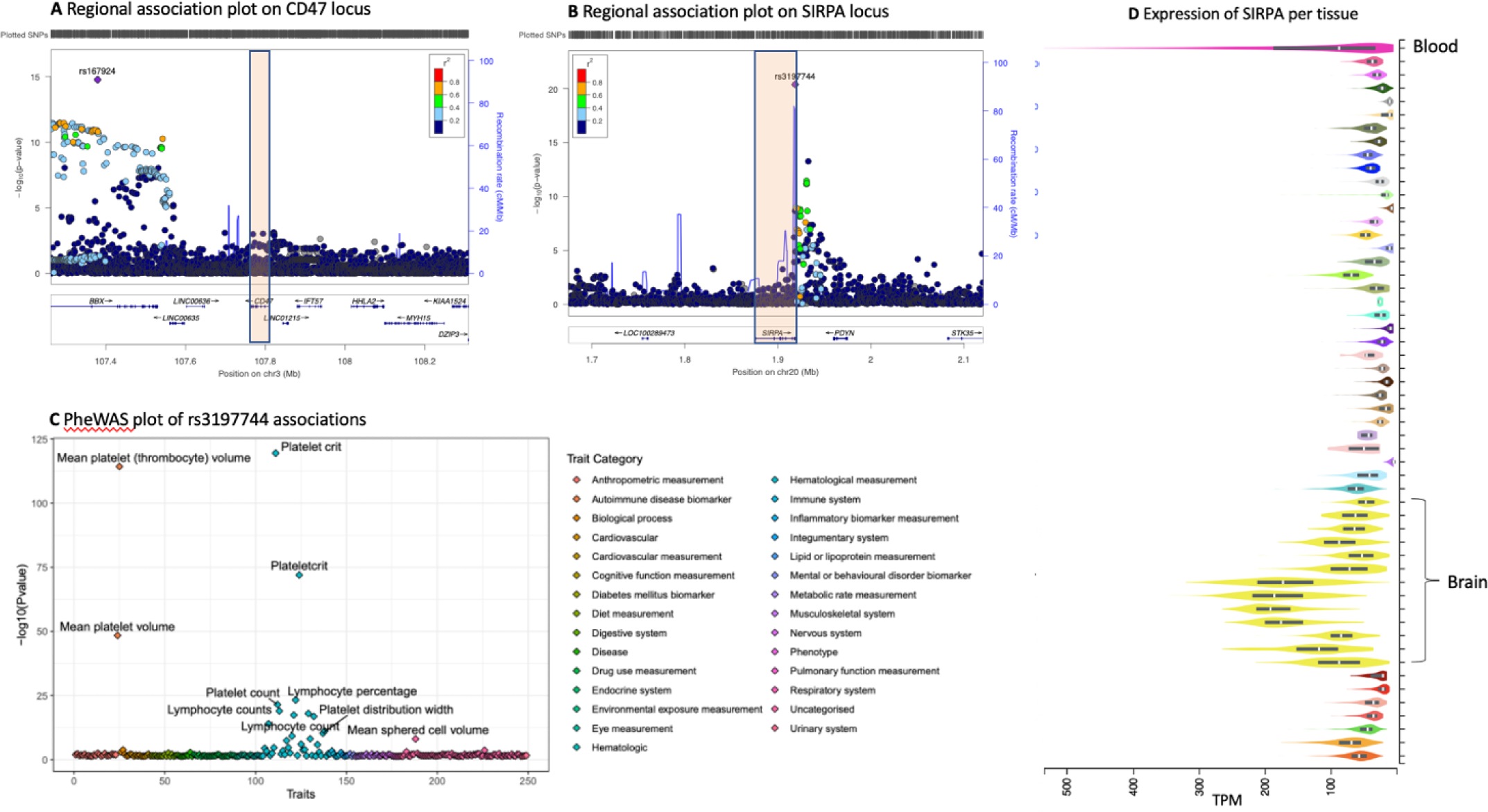
Variants at SIRPA are associated with platelet levels. (A) Variation at 3’ to CD47 at BBX locus associated with platelet levels. (B) Variant rs3197744 at SIRPA locus associates with amount of platelets. (C) Examining associations across currently available GWAS results (PheWAS) over 2500 independent studies at rs3197744 at SIRPA locus shows specific association with platelet and immune cell measurements. (D) Expression profile of SIRPA across human tissues shows highest expression in the brain and in blood.

We explored the phenotypic associations of these variants with stroke and coagulation traits, as well as other publicly available traits. We observed that the variants at *SIRPA* had specific associations with blood cell traits overall. The association with platelets was strongest, which has also been observed in previous GWAS ^29^(Fig. 3C, and Supplementary information). The expression profile of *SIRPA* RNA across human tissues observed the highest expression levels in whole blood and the brain (Fig. 3D), as would be expected from the high SIRPα protein expression observed on neurons and myeloid cells. Overall, these findings show a contribution from variants in the *SIRPA* 3’UTR and at *BBX-CD47* loci to platelet levels, in alignment with our functional studies.

### Genetic variation affecting structure, expression and disease associations on *SIRPA* locus

*SIRPA* is highly polymorphic, encompassing several intronic and coding variants. Genetic variation at the *SIRPA* locus has previously focused on exploring coding variation at the second exon, where missense variants can be divided into two common haplotypes that are referred to as V1 and V2 (Fig. 4). These variants are part of a haplotype from rs6132062 to rs6136377 that spans several regulatory elements from intron one to two and has the largest impact on *SIRPA* gene expression in blood ^30^. In addition, the regulatory variants affecting *SIRPA* gene expression are also in high linkage disequilibrium (LD, r^2^=1) with the V1 haplotype, so that V1 expression also correlates with the amount of overall *SIRPA* expression. In contrast, nearly all disease and trait associations with the *SIRPA* locus are located in the 3’UTR and are not in LD with the eQTL signal, or with V1 or V2 haplotypes. Consequently, the SNPs that are associated with platelet levels are also located at the 3’UTR of *SIRPA*.

**Figure 4.**
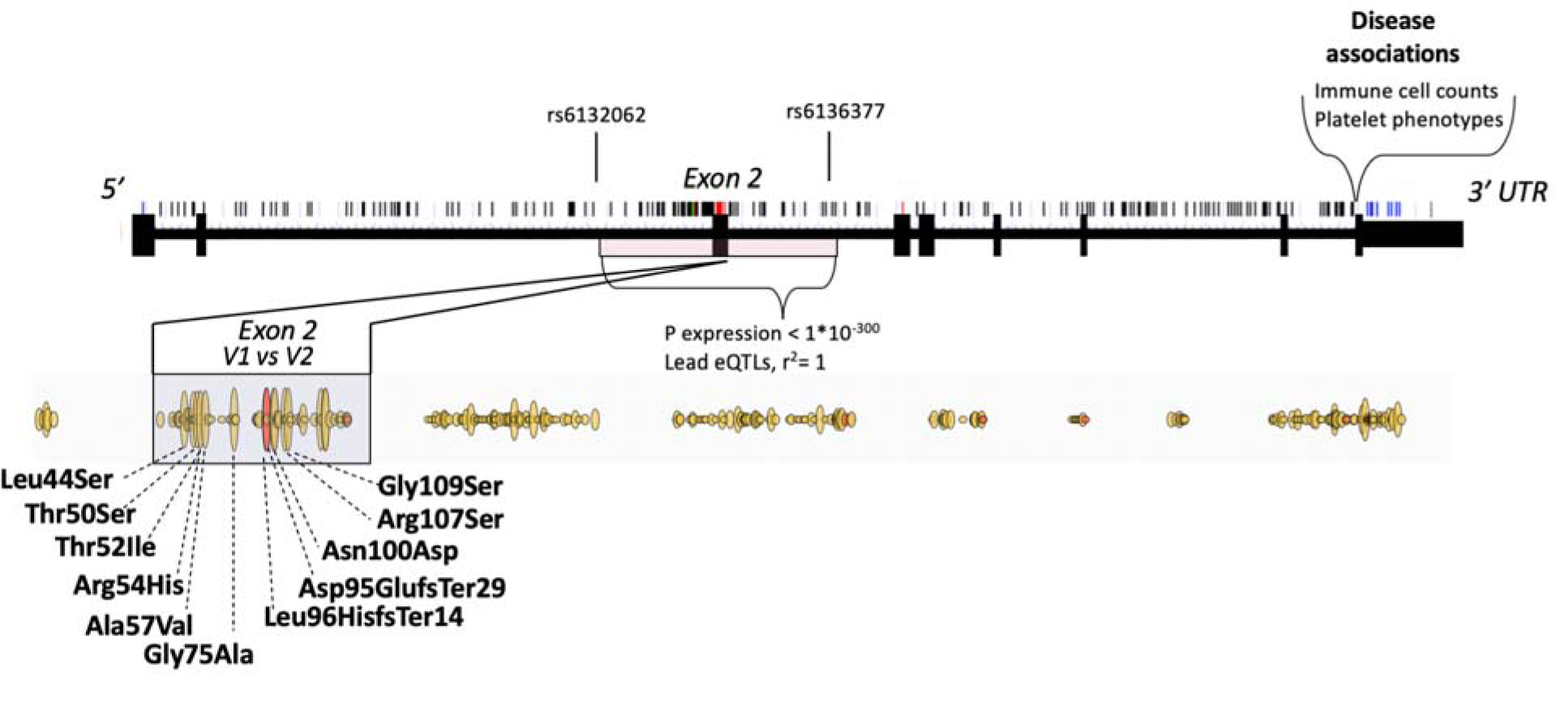
Schematic representation of genetic variation affecting structure, expression and disease associations on *SIRPA* locus. Single nucleotide polymorphisms are marked on top of the gene structure. Missense variants are highlighted in red as part of the gene structure and outlined yellow below the gene structure. Frameshift/insertions or deletions are highlighted with red as part of the missense variants. Haplotype from rs6132062 to rs6136377 encompassing exon 2 has the largest impact on SIRPA gene expression in blood. In addition, exon 2 contains missense variants that can be divided into two common haplotypes that are referred to as V1 and V2. V1 allele is shown first in the graph. V1 is in linkage disequilibrium (r2=1) with the SIRPA eQTLs (rs6132062 and rs6136377) having impact therefore both on expression and on coding sequence. Disease associations with SIRPA are located in the 3’UTR including the variants that associate with platelet levels.

### *SIRPA* expression and broader context of platelets in human disease

We expect these genetic variants to affect the number of platelets through one of two mechanisms. First, polymorphisms can lead to changes in amino acids and have a direct effect on protein function. Second, polymorphisms may occur at transcriptional regulatory regions, where they lead to changes in the amount or properties of the transcript, and thus affect the gene product or protein function. This is often cell type-specific and controlled by tissue-specific transcriptional regulators. Such differences can be explored through expression quantitative trait loci (eQTLs), location on active chromatin and tissue specificity, or transcription-wide analysis (TWAS). Indeed, when we examined *SIRPA* expression in humans and assessed its association with the amount of platelets using TWAS ^31^, we saw a significant positive correlation with *SIRPA* gene expression and the amount of platelets (Z=9.3, p= Best TWAS P=1.1e-19, Fig. 5A).

**Figure 5.**
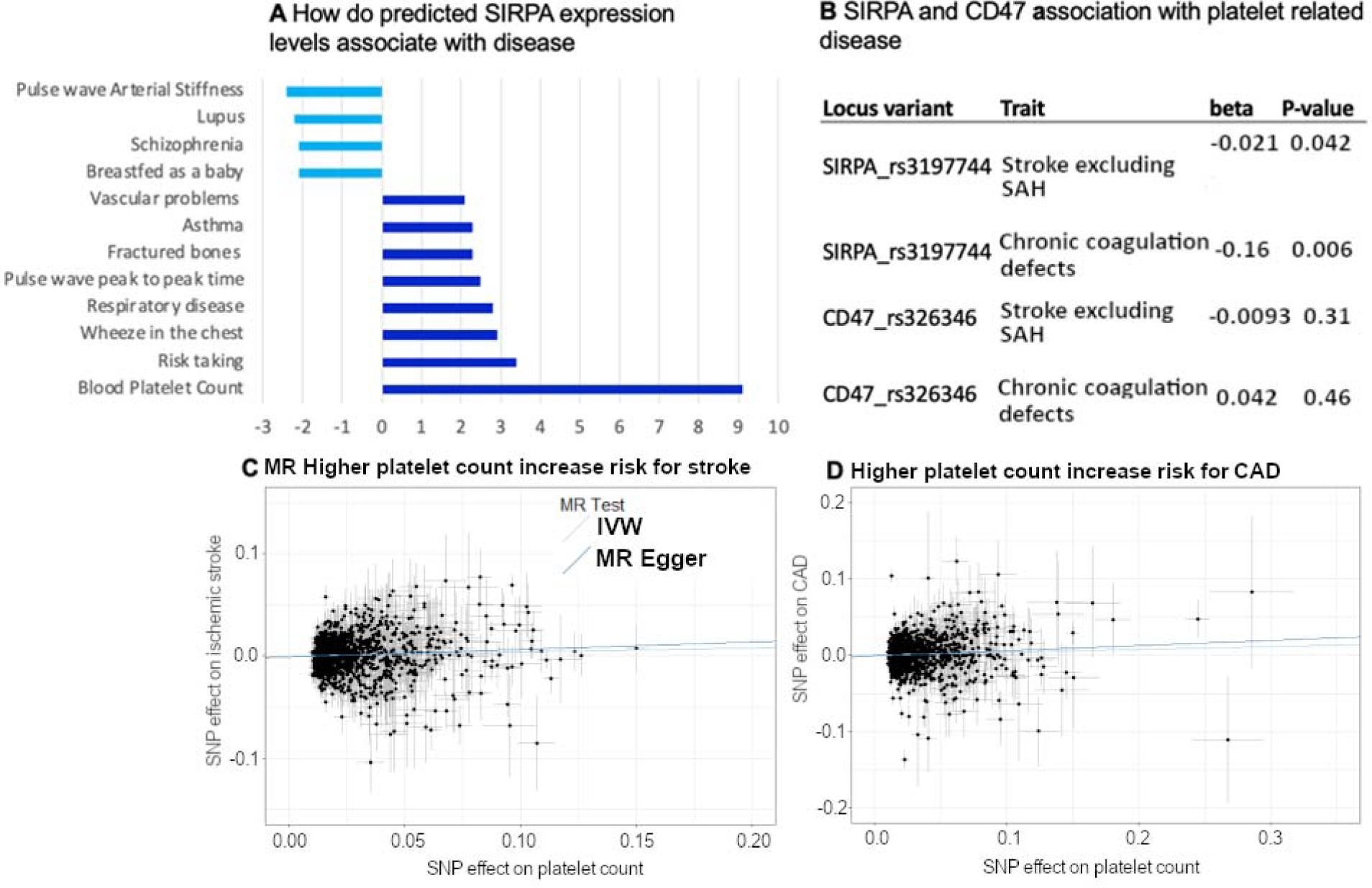
SIRPA expression and broader context of platelets in human disease. (A) TWAS analysis of SIRPA expression with human phenotypes. X-axis shows effect on expression by Z-statistics. (B) Association of leading SIRPA and CD47 SNPs with stroke excluding subarachnoid hemorrhage (SAH) and coagulation traits that have been implicated in platelet function. (C) Analysis of causality using Mendelian randomization from platelet levels to stroke, (D) and to coronary artery disease.

To understand the contribution from coding variants at the *SIRPA* locus, we explored the earlier computed burden analyses from the Genebass project (https://genebass.org/), which comprises analysis of variants in 281,852 individuals with exome sequence data from the UK Biobank. Burden analysis of missense variants at *SIRPA* showed an association of missense variants with a higher percentage of platelets in the blood as measured with plateletcrit (p = 0.014, Fig. 5A), but a nominal association with a subset of other platelet and reticulocyte traits (Supplementary information).

We then assessed whether *SIRPA* variants that contribute to platelet levels associate with other phenotypes known to be related to platelet levels. Indeed, the platelet-associated SNPs on *SIRPA* are nominally associated with coagulation and with ischemic stroke phenotypes in FinnGen (Fig. 5B). We also tested the causality behind higher platelet levels and ischemic stroke and coronary artery disease using genetic instruments and a Mendelian randomization approach. We observed a causal effect from higher platelet levels in higher risk for ischemic stroke and coronary artery disease (Fig. 5C and D).

### SIRPA and CD47 are inversely correlated, and are independently associated with increased platelet count in ET patients

We further evaluated the CD47-SIRPα axis in a second disease phenotype, essential thrombocythemia (ET), which is also associated with increased platelet counts ^32^. Here, we measured platelet transcriptomic expression of CD47 and SIRPα as a function of platelet counts in patients with ET (n=23) compared with that of healthy donors (n=11). CD47 and SIRPα platelet RNA expression levels were inversely correlated (Fig. 6A), with increasingly higher expression of SIRPA in ET patients compared to healthy donors. Therefore, as a function of platelet counts, CD47 expression levels were also inversely correlated with higher platelet counts in ET patients versus healthy donors, and vice versa for SIRPα expression levels (Fig. 6B and C).

**Figure 6.**
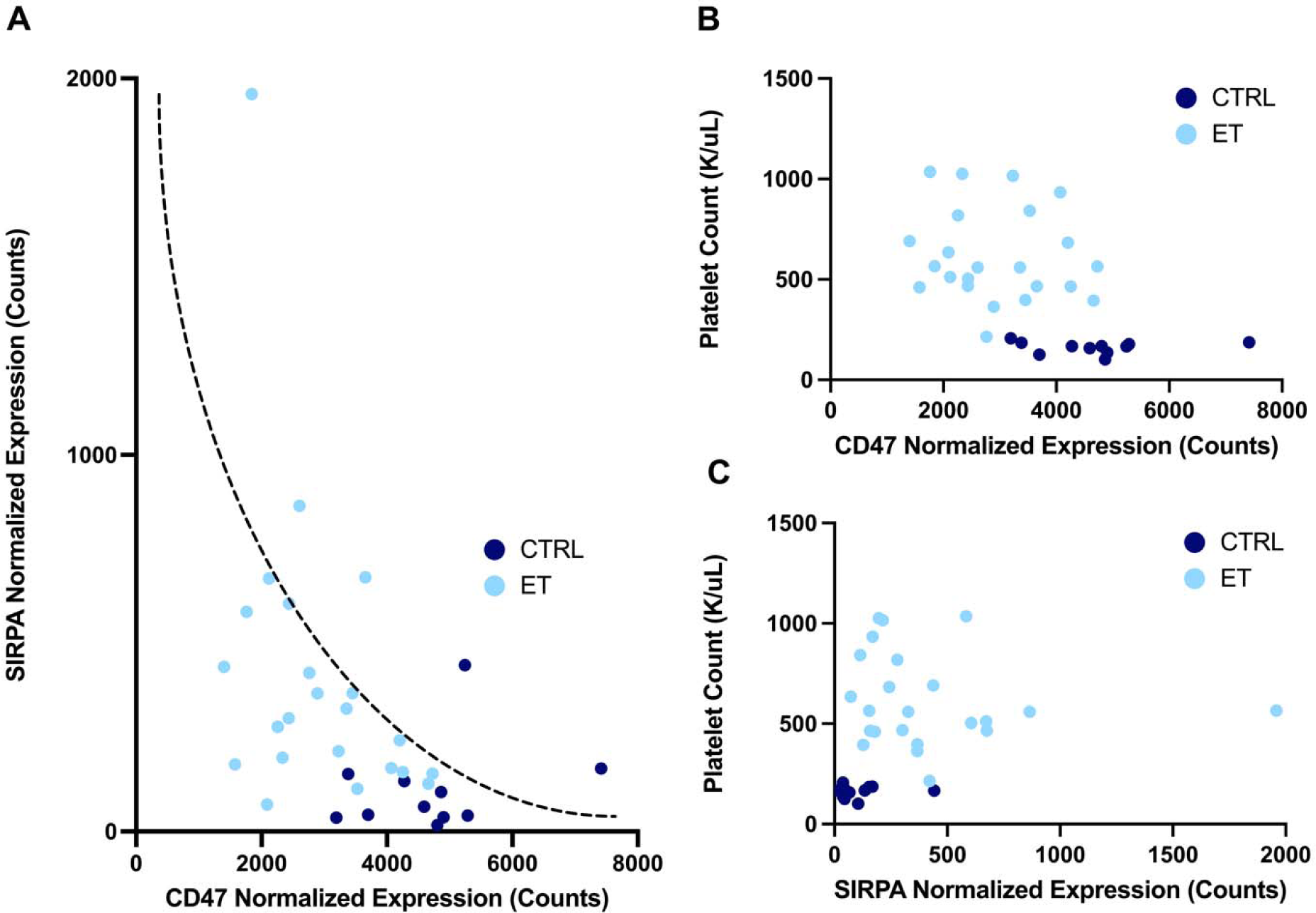
SIRPA and CD47 are inversely correlated, and are independently associated with platelet count in ET patients. (A) Correlation of platelet normalized gene expression between CD47 and SIRPA for patients with essential thrombocythemia and healthy donors. Closed circles navy or light blue indicate sample type respectively of healthy donors as controls (CTRL, n=11) and patients with essential thrombocythemia (ET, n=23). Inverse correlation is indicated by a dotted curve as a guide to the eye. Platelet gene expression as assessed by normalized counts of (B) SIRPA and (C) CD47 as a function of patient platelet counts (K/uL) at the time of specimen banking and platelet isolation. Closed circles navy or light blue indicate sample type respectively of healthy donors as controls (CTRL, n=11) and patients with essential thrombocythemia (ET, n=23).

## Discussion

We have shown that platelet homeostasis and clearance is regulated through the CD47-SIRPα axis and that therapeutic blockade to disrupt this interaction has a significant impact on platelet levels. Our findings summarize and validate previous observations from clinical studies, blockade assays from animal studies, and genetic evidence.

The CD47-SIRPα axis has an established role in platelet hematopoiesis, but the impact of the highly polymorphic *SIRPA* gene on population level heterogeneity of outcomes from blocking the CD47-SIRPa axis are not known. Yamao et al. (2002) ^33^ showed that modulation of SIRPα expression and signaling can increase platelet clearance and lead to insufficient compensatory megakaryocytopoiesis and platelet production. They used mice lacking the intracellular domain of SIRPα and showed increased splenic megakaryocytes and thrombocytopenia, similar to what we observed after blockade of CD47 in WT mice. CD47-deficient mice on a C57BL/6 background have also been shown to produce platelets with considerably shorter half-lives and consequently exhibit thrombocytopenia, with a dose-response effect between total CD47 knockouts, heterozygotes and wild-type mice ^9,34^. CD47^−/−^ platelets transferred into wild-type mice underwent increased and accelerated clearance, and blockade of SIRPα increases phagocytosis of wild-type platelets to the level of CD47^−/−^ platelets. Thus, deficiencies in either CD47 or SIRPα individually appear to be sufficient to drive up platelet clearance. This suggests that both, but not one or the other, are necessary for platelet survival and homeostasis. Our *in vivo* studies using CD47 blockade recapitulate these results, showing increased megakaryocytes in the spleen of mice treated with an antibody against the SIRPα-binding domain of CD47, and showing simultaneously reduced platelet count and increased reticulocyte count in the circulation. While the platelet numbers are reduced, individual platelets are hypertrophic, which raises the possibility of a compensatory increase not only in megakaryocyte number but also in the rate of megakaryocyte-platelet cleavage, resulting in larger platelet fragments. Clinical trials of anti-CD47 biologics have implied the existence of a similar mechanism in humans, though few studies exist that elucidate the details.

We hypothesized a connection between the observed clinical trial data and murine experiments with genome-wide association studies (GWAS), where we have identified genetic variants at the *SIRPA* locus 3’UTR that associate with platelet levels ^29^. To understand the association and possible underlying mechanisms, we performed a meta-analysis of publicly available cohorts. In our analysis, we first identified regulatory variants at intronic and untranslated regions of *SIRPA*. While the primary region showing ‘platelet effect’ is seen at the 3’UTR of *SIRPA*, our analysis also shows that higher *SIRPA* expression in humans is associated with higher platelet levels. Furthermore, the well-established V1-haplotype of non-synonymous genetic variants at exon 2 of *SIRPA* correlates with higher *SIRPA* expression. We identified three partially independent genetic mechanisms: a) 3’UTR association, b) intronic eQTLs, and c) non-synonymous coding variants that contribute at the *SIRPA* locus and translate to higher platelet levels.

Ultimately, the clinical relevance of platelet biology centers on individuals’ risk of thromboembolic events like stroke and CAD. Our analysis demonstrates a connection between SIRPA variants, platelet levels and risk of stroke and CAD. It is known that there exists a role for CD47 in atherosclerosis, where it has been shown that ApoE−/− mice on high fat diets develop atherosclerosis partly due to upregulation of CD47 on plaque smooth muscle cells ^35^. Anti-CD47 antibodies can help prevent atherosclerosis in monotherapy ^36^, and in combination with anti-TNFα can reduce preformed plaques, a process in which plaque macrophages remove necrotic cores by efferocytosis as well as pathogenic clonal smooth muscle cells by programmed cell removal (PrCR) ^36,37^. Since the CD47-SIRPα axis is known to contribute to atherosclerosis, this prompts the idea that genetic polymorphisms within the axis can impact the potential of axis modulation in stroke and CAD.

To further demonstrate the link between CD47 and SIRPα expression levels and platelet count. We analyzed the CD47-SIRPα axis in the context of ET. ET patients with increased platelet levels exhibit elevated SIRPα platelet RNA expression levels in comparison to healthy patients, and the SIRPα expression levels are inversely correlated to CD47 levels. Disruption of the CD47-SIRPα axis seems to directly impact platelet levels, suggesting that genetic variation within the axis may have a similar effect.

Taken together, our findings bring up a novel aspect of CD47 blockade in the context of genetic polymorphisms of *SIRPA* that can impact platelet biology. Future studies should investigate the extent to which these polymorphisms could potentially help identify patients at greater risk of developing platelet-related diseases, including thrombocytopenia, ET, CAD, and stroke, with treatments that interfere with this signaling pathway and should probe further the functional impact of these differences in platelet levels and characteristics.

## Supporting information

FinnGen Ethics Statement

FinnGen Banner

## Acknowledgments

We want to acknowledge the participants and investigators of the FinnGen study. The authors wish to thank members of the Weissman Lab at the Institute for Stem Cell Biology & Regenerative Medicine at Stanford University School of Medicine and Histowiz for their processing of our histological samples. We also thank Dr. Jason Gotlib at Stanford University, MPN ET patients at the Stanford Cancer Institute, and the healthy donors at the Stanford Blood Center for their contribution to the platelet transcriptome research that enabled further study of select candidate genes in this work.

## Authorship Contributions

M.C.T performed experiments, wrote and edited the manuscript, and supervised the research. H.M.O analyzed data, wrote and edited the manuscript, and supervised the research. A.K. supervised the research (in particular platelet RNA seq experiments), analyzed the data and edited the manuscript. M.S, Y.Y.Y and P.S.H performed experiments, wrote, and edited the manuscript. A.S. analyzed data and wrote and edited the manuscript. M.B analyzed data and wrote and edited the manuscript. E.G. analyzed data. N.S. analyzed data, wrote and edited the manuscript, and supervised the research, I.L.W supervised the research. FinnGen provided and analyzed data.

