## Supplementary material for "SIRPα controls CD47-dependent platelet clearance in mice and humans": FinnGen Banner

**Expanded FinnGen contributor list.** List of primary contributors involved with FinnGen R9.

**Steering Committee**

Aarno Palotie	Institute for Molecular Medicine Finland, HiLIFE, University of Helsinki, Finland

Mark Daly	Institute for Molecular Medicine Finland, HiLIFE, University of Helsinki, Finland

**Pharmaceutical companies**

Bridget Riley-Gills	Abbvie, Chicago, IL, United States

Howard Jacob	Abbvie, Chicago, IL, United States

Dirk Paul	Astra Zeneca, Cambridge, United Kingdom

Heiko Runz	Biogen, Cambridge, MA, United States

Sally John	Biogen, Cambridge, MA, United States

George Okafo	Boehringer Ingelheim, Ingelheim am Rhein, Germany

Nathan Lawless	Boehringer Ingelheim, Ingelheim am Rhein, Germany

Robert Plenge	Celgene, Summit, NJ, United States/Bristol Myers Squibb, New York, NY, United States

Joseph Maranville	Celgene, Summit, NJ, United States/Bristol Myers Squibb, New York, NY, United States

Mark McCarthy	Genentech, San Francisco, CA, United States

Julie Hunkapiller	Genentech, San Francisco, CA, United States

Meg Ehm	GlaxoSmithKline, Brentford, United Kingdom

Kirsi Auro	GlaxoSmithKline, Brentford, United Kingdom

Simonne Longerich	Merck, Kenilworth, NJ, United States

Caroline Fox	Merck, Kenilworth, NJ, United States

Anders Mälarstig	Pfizer, New York, NY, United States

Katherine Klinger	Sanofi, Paris, France

Deepak Raipal	Sanofi, Paris, France

Eric Green	Maze Therapeutics, San Francisco, CA, United States

Robert Graham	Maze Therapeutics, San Francisco, CA, United States

Robert Yang	Janssen Biotech, Beerse, Belgium

Chris O´Donnell	Novartis, Basel, Switzerland

**University of Helsinki & Biobanks**

Tomi Mäkelä	HiLIFE, University of Helsinki, Finland, Finland

Jaakko Kaprio	Institute for Molecular Medicine Finland, HiLIFE, Helsinki, Finland, Finland

Petri Virolainen	Auria Biobank / University of Turku / Hospital District of Southwest Finland, Turku, Finland

Antti Hakanen	Auria Biobank / University of Turku / Hospital District of Southwest Finland, Turku, Finland

Terhi Kilpi	THL Biobank / The National Institute of Health and Welfare Helsinki, Finland

Markus Perola	THL Biobank / The National Institute of Health and Welfare Helsinki, Finland

Jukka Partanen	Finnish Red Cross Blood Service / Finnish Hematology Registry and Clinical Biobank, Helsinki, Finland

Anne Pitkäranta	Helsinki Biobank / Helsinki University and Hospital District of Helsinki and Uusimaa, Helsinki

Juhani Junttila	Northern Finland Biobank Borealis / University of Oulu / Northern Ostrobothnia Hospital District, Oulu, Finland

Raisa Serpi	Northern Finland Biobank Borealis / University of Oulu / Northern Ostrobothnia Hospital District, Oulu, Finland

Tarja Laitinen	Finnish Clinical Biobank Tampere **/** University of Tampere / Pirkanmaa Hospital District, Tampere, Finland

Veli-Matti Kosma	Biobank of Eastern Finland / University of Eastern Finland / Northern Savo Hospital District, Kuopio, Finland

Jari Laukkanen	Central Finland Biobank / University of Jyväskylä / Central Finland Health Care District, Jyväskylä, Finland

Marco Hautalahti	FINBB - Finnish biobank cooperative

**Other Experts/ Non-Voting Members**

Outi Tuovila	Business Finland, Helsinki, Finland

Raimo Pakkanen	Business Finland, Helsinki, Finland

**Scientific Committee**

**Pharmaceutical companies**

Jeffrey Waring	Abbvie, Chicago, IL, United States

Bridget Riley-Gillis	Abbvie, Chicago, IL, United States

Ioanna Tachmazidou	Astra Zeneca, Cambridge, United Kingdom

Chia-Yen Chen	Biogen, Cambridge, MA, United States

Heiko Runz	Biogen, Cambridge, MA, United States

Zhihao Ding	Boehringer Ingelheim, Ingelheim am Rhein, Germany

Marc Jung	Boehringer Ingelheim, Ingelheim am Rhein, Germany

Shameek Biswas	Celgene, Summit, NJ, United States/Bristol Myers Squibb, New York, NY, United States

Rion Pendergrass	Genentech, San Francisco, CA, United States

Julie Hunkapiller	Genentech, San Francisco, CA, United States

Meg Ehm	GlaxoSmithKline, Brentford, United Kingdom

David Pulford	GlaxoSmithKline, Brentford, United Kingdom

Neha Raghavan	Merck, Kenilworth, NJ, United States

Adriana Huertas-Vazquez	Merck, Kenilworth, NJ, United States

Jae-Hoon Sul	Merck, Kenilworth, NJ, United States

Anders Mälarstig	Pfizer, New York, NY, United States

Xinli Hu	Pfizer, New York, NY, United States

Katherine Klinger	Sanofi, Paris, France

Matthias Gossel	Sanofi, Paris, France

Robert Graham	Maze Therapeutics, San Francisco, CA, United States

Eric Green	Maze Therapeutics, San Francisco, CA, United States

Sahar Mozaffari	Maze Therapeutics, San Francisco, CA, United States

Dawn Waterworth	Janssen Research & Development, LLC, Spring House, PA, United States

Nicole Renaud	Novartis, Basel, Switzerland

Ma´en Obeidat	Novartis, Basel, Switzerland

**University of Helsinki & Biobanks**

Samuli Ripatti	Institute for Molecular Medicine Finland, HiLIFE, Helsinki, Finland

Johanna Schleutker	Auria Biobank / Univ. of Turku / Hospital District of Southwest Finland, Turku, Finland

Markus Perola	THL Biobank / The National Institute of Health and Welfare Helsinki, Finland

Mikko Arvas	Finnish Red Cross Blood Service / Finnish Hematology Registry and Clinical Biobank, Helsinki, Finland

Olli Carpén	Helsinki Biobank / Helsinki University and Hospital District of Helsinki and Uusimaa, Helsinki

Reetta Hinttala	Northern Finland Biobank Borealis / University of Oulu / Northern Ostrobothnia Hospital District, Oulu, Finland

Johannes Kettunen	Northern Finland Biobank Borealis / University of Oulu / Northern Ostrobothnia Hospital District, Oulu, Finland

Arto Mannermaa	Biobank of Eastern Finland / University of Eastern Finland / Northern Savo Hospital District, Kuopio, Finland

Katriina Aalto-Setälä	Finnish Clinical Biobank Tampere **/** University of Tampere / Pirkanmaa Hospital District, Tampere, Finland

Mika Kähönen	Finnish Clinical Biobank Tampere **/** University of Tampere / Pirkanmaa Hospital District, Tampere, Finland

Jari Laukkanen	Central Finland Biobank / University of Jyväskylä / Central Finland Health Care District, Jyväskylä, Finland

Johanna Mäkelä	FINBB - Finnish biobank cooperative

**Clinical Groups**

**Neurology Group**

Reetta Kälviäinen	Northern Savo Hospital District, Kuopio, Finland

Valtteri Julkunen	Northern Savo Hospital District, Kuopio, Finland

Hilkka Soininen	Northern Savo Hospital District, Kuopio, Finland

Anne Remes	Northern Ostrobothnia Hospital District, Oulu, Finland

Mikko Hiltunen	Northern Savo Hospital District, Kuopio, Finland

Jukka Peltola	Pirkanmaa Hospital District, Tampere, Finland

Minna Raivio	Hospital District of Helsinki and Uusimaa, Helsinki, Finland

Pentti Tienari	Hospital District of Helsinki and Uusimaa, Helsinki, Finland

Juha Rinne	Hospital District of Southwest Finland, Turku, Finland

Roosa Kallionpää	Hospital District of Southwest Finland, Turku, Finland

Juulia Partanen	Institute for Molecular Medicine Finland, HiLIFE, University of Helsinki, Finland

Ali Abbasi	Abbvie, Chicago, IL, United States

Adam Ziemann	Abbvie, Chicago, IL, United States

Jeffrey Waring	Abbvie, Chicago, IL, United States

Nizar Smaoui	Abbvie, Chicago, IL, United States

Anne Lehtonen	Abbvie, Chicago, IL, United States

Susan Eaton	Biogen, Cambridge, MA, United States

Heiko Runz	Biogen, Cambridge, MA, United States

Sanni Lahdenperä	Biogen, Cambridge, MA, United States

Janet van Adelsberg	Celgene, Summit, NJ, United States/ Bristol Myers Squibb, New York, NY, United States

Shameek Biswas	Celgene, Summit, NJ, United States/ Bristol Myers Squibb, New York, NY, United States

Julie Hunkapiller	Genentech, San Francisco, CA, United States

Natalie Bowers	Genentech, San Francisco, CA, United States

Edmond Teng	Genentech, San Francisco, CA, United States

Rion Pendergrass	Genentech, San Francisco, CA, United States

Fanli Xu	GlaxoSmithKline, Brentford, United Kingdom

David Pulford	GlaxoSmithKline, Brentford, United Kingdom

Kirsi Auro	GlaxoSmithKline, Brentford, United Kingdom

Laura Addis	GlaxoSmithKline, Brentford, United Kingdom

John Eicher	GlaxoSmithKline, Brentford, United Kingdom

Qingqin S Li	Janssen Research & Development, LLC, Titusville, NJ 08560, United States

Karen He	Janssen Research & Development, LLC, Spring House, PA, United States

Ekaterina Khramtsova	Janssen Research & Development, LLC, Spring House, PA, United States

Neha Raghavan	Merck, Kenilworth, NJ, United States

Kari Linden	Pfizer, New York, NY, United States

**Gastroenterology Group**

Martti Färkkilä	Hospital District of Helsinki and Uusimaa, Helsinki, Finland

Jukka Koskela	Hospital District of Helsinki and Uusimaa, Helsinki, Finland

Sampsa Pikkarainen	Hospital District of Helsinki and Uusimaa, Helsinki, Finland

Airi Jussila	Pirkanmaa Hospital District, Tampere, Finland

Katri Kaukinen	Pirkanmaa Hospital District, Tampere, Finland

Timo Blomster	Northern Ostrobothnia Hospital District, Oulu, Finland

Mikko Kiviniemi	Northern Savo Hospital District, Kuopio, Finland

Markku Voutilainen	Hospital District of Southwest Finland, Turku, Finland

Mark Daly	Institute for Molecular Medicine Finland, HiLIFE, University of Helsinki, Finland

Ali Abbasi	Abbvie, Chicago, IL, United States

Graham Heap	Abbvie, Chicago, IL, United States

Jeffrey Waring	Abbvie, Chicago, IL, United States

Nizar Smaoui	Abbvie, Chicago, IL, United States

Fedik Rahimov	Abbvie, Chicago, IL, United States

Anne Lehtonen	Abbvie, Chicago, IL, United States

Keith Usiskin	Celgene, Summit, NJ, United States/ Bristol Myers Squibb, New York, NY, United States

Tim Lu	Genentech, San Francisco, CA, United States

Natalie Bowers	Genentech, San Francisco, CA, United States

Rion Pendergrass	Genentech, San Francisco, CA, United States

Linda McCarthy	GlaxoSmithKline, Brentford, United Kingdom

Amy Hart	Janssen Research & Development, LLC, Spring House, PA, United States

Meijian Guan	Janssen Research & Development, LLC, Spring House, PA, United States

Jason Miller	Merck, Kenilworth, NJ, United States

Kirsi Kalpala	Pfizer, New York, NY, United States

Melissa Miller	Pfizer, New York, NY, United States

Xinli Hu	Pfizer, New York, NY, United States

**Rheumatology Group**

Kari Eklund	Hospital District of Helsinki and Uusimaa, Helsinki, Finland

Antti Palomäki	Hospital District of Southwest Finland, Turku, Finland

Pia Isomäki	Pirkanmaa Hospital District, Tampere, Finland

Laura Pirilä	Hospital District of Southwest Finland, Turku, Finland

Oili Kaipiainen-Seppänen	Northern Savo Hospital District, Kuopio, Finland

Johanna Huhtakangas	Northern Ostrobothnia Hospital District, Oulu, Finland

Nina Mars	Institute for Molecular Medicine Finland, HiLIFE, Helsinki, Finland

Ali Abbasi	Abbvie, Chicago, IL, United States

Jeffrey Waring	Abbvie, Chicago, IL, United States

Fedik Rahimov	Abbvie, Chicago, IL, United States

Apinya Lertratanakul	Abbvie, Chicago, IL, United States

Nizar Smaoui	Abbvie, Chicago, IL, United States

Anne Lehtonen	Abbvie, Chicago, IL, United States

David Close	Astra Zeneca, Cambridge, United Kingdom

Marla Hochfeld	Celgene, Summit, NJ, United States/ Bristol Myers Squibb, New York, NY, United States

Natalie Bowers	Genentech, San Francisco, CA, United States

Rion Pendergrass	Genentech, San Francisco, CA, United States

Jorge Esparza Gordillo	GlaxoSmithKline, Brentford, United Kingdom

Kirsi Auro	GlaxoSmithKline, Brentford, United Kingdom

Dawn Waterworth	Janssen Research & Development, LLC, Spring House, PA, United States

Fabiana Farias	Merck, Kenilworth, NJ, United States

Kirsi Kalpala	Pfizer, New York, NY, United States

Nan Bing	Pfizer, New York, NY, United States

Xinli Hu	Pfizer, New York, NY, United States

**Pulmonology Group**

Tarja Laitinen	Pirkanmaa Hospital District, Tampere, Finland

Margit Pelkonen	Northern Savo Hospital District, Kuopio, Finland

Paula Kauppi	Hospital District of Helsinki and Uusimaa, Helsinki, Finland

Hannu Kankaanranta	University of Gothenburg, Gothenburg, Sweden/ Seinäjoki Central Hospital, Seinäjoki, Finland/ Tampere University, Tampere, Finland

Terttu Harju	Northern Ostrobothnia Hospital District, Oulu, Finland

Riitta Lahesmaa	Hospital District of Southwest Finland, Turku, Finland

Nizar Smaoui	Abbvie, Chicago, IL, United States

Alex Mackay	Astra Zeneca, Cambridge, United Kingdom

Glenda Lassi	Astra Zeneca, Cambridge, United Kingdom

Susan Eaton	Biogen, Cambridge, MA, United States

Steven Greenberg	Celgene, Summit, NJ, United States/ Bristol Myers Squibb, New York, NY, United States

Hubert Chen	Genentech, San Francisco, CA, United States

Rion Pendergrass	Genentech, San Francisco, CA, United States

Natalie Bowers	Genentech, San Francisco, CA, United States

Joanna Betts	GlaxoSmithKline, Brentford, United Kingdom

Kirsi Auro	GlaxoSmithKline, Brentford, United Kingdom

Rajashree Mishra	GlaxoSmithKline, Brentford, United Kingdom

Majd Mouded	Novartis, Basel, Switzerland

Debby Ngo	Novartis, Basel, Switzerland

**Cardiometabolic Diseases Group**

Teemu Niiranen	The National Institute of Health and Welfare Helsinki, Finland

Felix Vaura	The National Institute of Health and Welfare Helsinki, Finland

Veikko Salomaa	The National Institute of Health and Welfare Helsinki, Finland

Kaj Metsärinne	Hospital District of Southwest Finland, Turku, Finland

Jenni Aittokallio	Hospital District of Southwest Finland, Turku, Finland

Mika Kähönen	Pirkanmaa Hospital District, Tampere, Finland

Jussi Hernesniemi	Pirkanmaa Hospital District, Tampere, Finland

Juhani Junttila	Northern Ostrobothnia Hospital District, Oulu, Finland

Markku Laakso	Northern Savo Hospital District, Kuopio, Finland

Jussi Pihlajamäki	Northern Savo Hospital District, Kuopio, Finland

Daniel Gordin	Hospital District of Helsinki and Uusimaa, Helsinki, Finland

Juha Sinisalo	Hospital District of Helsinki and Uusimaa, Helsinki, Finland

Marja-Riitta Taskinen	Hospital District of Helsinki and Uusimaa, Helsinki, Finland

Tiinamaija Tuomi	Hospital District of Helsinki and Uusimaa, Helsinki, Finland

Timo Hiltunen	Hospital District of Helsinki and Uusimaa, Helsinki, Finland

Jari Laukkanen	Central Finland Health Care District, Jyväskylä, Finland

Amanda Elliott	Institute for Molecular Medicine Finland, HiLIFE, University of Helsinki, Finland / Broad Institute, Cambridge, MA, United States

Mary Pat Reeve	Institute for Molecular Medicine Finland, HiLIFE, University of Helsinki, Finland

Sanni Ruotsalainen	Institute for Molecular Medicine Finland, HiLIFE, University of Helsinki, Finland

Benjamin Challis	Astra Zeneca, Cambridge, United Kingdom

Dirk Paul	Astra Zeneca, Cambridge, United Kingdom

Keith Usiskin	Celgene, Summit, NJ, United States/ Bristol Myers Squibb, New York, NY, United States

Julie Hunkapiller	Genentech, San Francisco, CA, United States

Natalie Bowers	Genentech, San Francisco, CA, United States

Rion Pendergrass	Genentech, San Francisco, CA, United States

Audrey Chu	GlaxoSmithKline, Brentford, United Kingdom

Kirsi Auro	GlaxoSmithKline, Brentford, United Kingdom

Dermot Reilly	Janssen Research & Development, LLC, Boston, MA, United States

Mike Mendelson	Novartis, Boston, MA, United States

Jaakko Parkkinen	Pfizer, New York, NY, United States

Melissa Miller	Pfizer, New York, NY, United States

**Oncology Group**

Tuomo Meretoja	Hospital District of Helsinki and Uusimaa, Helsinki, Finland

Heikki Joensuu	Hospital District of Helsinki and Uusimaa, Helsinki, Finland

Olli Carpén	Hospital District of Helsinki and Uusimaa, Helsinki, Finland

Lauri Aaltonen	Hospital District of Helsinki and Uusimaa, Helsinki, Finland

Johanna Mattson	Hospital District of Helsinki and Uusimaa, Helsinki, Finland

Eveliina Salminen	Hospital District of Helsinki and Uusimaa, Helsinki, Finland

Annika Auranen	Pirkanmaa Hospital District , Tampere, Finland

Peeter Karihtala	Northern Ostrobothnia Hospital District, Oulu, Finland

Päivi Auvinen	Northern Savo Hospital District, Kuopio, Finland

Klaus Elenius	Hospital District of Southwest Finland, Turku, Finland

Johanna Schleutker	Hospital District of Southwest Finland, Turku, Finland

Esa Pitkänen	Institute for Molecular Medicine Finland, HiLIFE, University of Helsinki, Finland

Nina Mars	Institute for Molecular Medicine Finland, HiLIFE, University of Helsinki, Finland

Mark Daly	Institute for Molecular Medicine Finland, HiLIFE, University of Helsinki, Finland

Relja Popovic	Abbvie, Chicago, IL, United States

Jeffrey Waring	Abbvie, Chicago, IL, United States

Bridget Riley-Gillis	Abbvie, Chicago, IL, United States

Anne Lehtonen	Abbvie, Chicago, IL, United States

Jennifer Schutzman	Genentech, San Francisco, CA, United States

Julie Hunkapiller	Genentech, San Francisco, CA, United States

Natalie Bowers	Genentech, San Francisco, CA, United States

Rion Pendergrass	Genentech, San Francisco, CA, United States

Diptee Kulkarni	GlaxoSmithKline, Brentford, United Kingdom

Kirsi Auro	GlaxoSmithKline, Brentford, United Kingdom

Alessandro Porello	Janssen Research & Development, LLC, Spring House, PA, United States

Andrey Loboda	Merck, Kenilworth, NJ, United States

Heli Lehtonen	Pfizer, New York, NY, United States

Stefan McDonough	Pfizer, New York, NY, United States

Marika Crohns	Sanofi, Paris, France

Sauli Vuoti	Sanofi, Paris, France

**Opthalmology Group**

Kai Kaarniranta	Northern Savo Hospital District, Kuopio, Finland

Joni A Turunen	Hospital District of Helsinki and Uusimaa, Helsinki, Finland

Terhi Ollila	Hospital District of Helsinki and Uusimaa, Helsinki, Finland

Hannu Uusitalo	Pirkanmaa Hospital District, Tampere, Finland

Juha Karjalainen	Institute for Molecular Medicine Finland, HiLIFE, University of Helsinki, Finland

Esa Pitkänen	Institute for Molecular Medicine Finland, HiLIFE, University of Helsinki, Finland

Mengzhen Liu	Abbvie, Chicago, IL, United States

Heiko Runz	Biogen, Cambridge, MA, United States

Stephanie Loomis	Biogen, Cambridge, MA, United States

Erich Strauss	Genentech, San Francisco, CA, United States

Natalie Bowers	Genentech, San Francisco, CA, United States

Hao Chen	Genentech, San Francisco, CA, United States

Rion Pendergrass	Genentech, San Francisco, CA, United States

**Dermatology Group**

Kaisa Tasanen	Northern Ostrobothnia Hospital District, Oulu, Finland

Laura Huilaja	Northern Ostrobothnia Hospital District, Oulu, Finland

Katariina Hannula-Jouppi	Hospital District of Helsinki and Uusimaa, Helsinki, Finland

Teea Salmi	Pirkanmaa Hospital District, Tampere, Finland

Sirkku Peltonen	Hospital District of Southwest Finland, Turku, Finland

Leena Koulu	Hospital District of Southwest Finland, Turku, Finland

Nizar Smaoui	Abbvie, Chicago, IL, United States

Fedik Rahimov	Abbvie, Chicago, IL, United States

Anne Lehtonen	Abbvie, Chicago, IL, United States

David Choy	Genentech, San Francisco, CA, United States

Rion Pendergrass	Genentech, San Francisco, CA, United States

Dawn Waterworth	Janssen Research & Development, LLC, Spring House, PA, United States

Kirsi Kalpala	Pfizer, New York, NY, United States

Ying Wu	Pfizer, New York, NY, United States

**Odontology Group**

Pirkko Pussinen	Hospital District of Helsinki and Uusimaa, Helsinki, Finland

Aino Salminen	Hospital District of Helsinki and Uusimaa, Helsinki, Finland

Tuula Salo	Hospital District of Helsinki and Uusimaa, Helsinki, Finland

David Rice	Hospital District of Helsinki and Uusimaa, Helsinki, Finland

Pekka Nieminen	Hospital District of Helsinki and Uusimaa, Helsinki, Finland

Ulla Palotie	Hospital District of Helsinki and Uusimaa, Helsinki, Finland

Maria Siponen	Northern Savo Hospital District, Kuopio, Finland

Liisa Suominen	Northern Savo Hospital District, Kuopio, Finland

Päivi Mäntylä	Northern Savo Hospital District, Kuopio, Finland

Ulvi Gursoy	Hospital District of Southwest Finland, Turku, Finland

Vuokko Anttonen	Northern Ostrobothnia Hospital District, Oulu, Finland

Kirsi Sipilä	Northern Ostrobothnia Hospital District, Oulu, Finland

Rion Pendergrass	Genentech, San Francisco, CA, United States

**Women’s Health and Reproduction Group**

Hannele Laivuori	Institute for Molecular Medicine Finland, HiLIFE, University of Helsinki, Finland

Venla Kurra	Pirkanmaa Hospital District, Tampere, Finland

Laura Kotaniemi-Talonen	Pirkanmaa Hospital District, Tampere, Finland

Oskari Heikinheimo	Hospital District of Helsinki and Uusimaa, Helsinki, Finland

Ilkka Kalliala	Hospital District of Helsinki and Uusimaa, Helsinki, Finland

Lauri Aaltonen	Hospital District of Helsinki and Uusimaa, Helsinki, Finland

Varpu Jokimaa	Hospital District of Southwest Finland, Turku, Finland

Johannes Kettunen	Northern Ostrobothnia Hospital District, Oulu, Finland

Marja Vääräsmäki	Northern Ostrobothnia Hospital District, Oulu, Finland

Outi Uimari	Northern Ostrobothnia Hospital District, Oulu, Finland

Laure Morin-Papunen	Northern Ostrobothnia Hospital District, Oulu, Finland

Maarit Niinimäki	Northern Ostrobothnia Hospital District, Oulu, Finland

Terhi Piltonen	Northern Ostrobothnia Hospital District, Oulu, Finland

Katja Kivinen	Institute for Molecular Medicine Finland, HiLIFE, University of Helsinki, Finland

Elisabeth Widen	Institute for Molecular Medicine Finland, HiLIFE, University of Helsinki, Finland

Taru Tukiainen	Institute for Molecular Medicine Finland, HiLIFE, University of Helsinki, Finland

Mary Pat Reeve	Institute for Molecular Medicine Finland, HiLIFE, University of Helsinki, Finland

Mark Daly	Institute for Molecular Medicine Finland, HiLIFE, University of Helsinki, Finland

Liu Aoxing	Institute for Molecular Medicine Finland, HiLIFE, University of Helsinki, Finland

Andrea Ganna	Institute for Molecular Medicine Finland, HiLIFE, University of Helsinki, Finland

Niko Välimäki	University of Helsinki, Helsinki, Finland

Eija Laakkonen	University of Jyväskylä, Jyväskylä, Finland

Jaakko Tyrmi	University of Oulu, Oulu, Finland / University of Tampere, Tampere, Finland

Heidi Silven	University of Oulu, Oulu, Finland

Eeva Slitz	University of Oulu, Oulu, Finland

Riikka Arffman	University of Oulu, Oulu, Finland

Susanna Savukoski	University of Oulu, Oulu, Finland

Triin Laisk	Estonian biobank, Tartu, Estonia

Natalia Pujol	Estonian biobank, Tartu, Estonia

Bridget Riley-Gillis	Abbvie, Chicago, IL, United States

Mengzhen Liu	Abbvie, Chicago, IL, United States

Rion Pendergrass	Genentech, San Francisco, CA, United States

Janet Kumar	GlaxoSmithKline, Brentford, United Kingdom

Kirsi Auro	GlaxoSmithKline, Brentford, United Kingdom

**FinnGen Analysis working group**

Bridget Riley-Gillis	Abbvie, Chicago, IL, United States

Reza Hammond	Abbvie, Chicago, IL, United States

Fedik Rahimov	Abbvie, Chicago, IL, United States

Sabah Kadri	Abbvie, Chicago, IL, United States

Mengzhen Liu	Abbvie, Chicago, IL, United States

Slavé Petrovski	Astra Zeneca, Cambridge, United Kingdom

Eleonor Wigmore	Astra Zeneca, Cambridge, United Kingdom

Adele Mitchell	Biogen, Cambridge, MA, United States

Benjamin Sun	Biogen, Cambridge, MA, United States

Ellen Tsai	Biogen, Cambridge, MA, United States

Denis Baird	Biogen, Cambridge, MA, United States

Paola Bronson	Biogen, Cambridge, MA, United States

Ruoyu Tian	Biogen, Cambridge, MA, United States

Stephanie Loomis	Biogen, Cambridge, MA, United States

Yunfeng Huang	Biogen, Cambridge, MA, United States

Till Andlauer	Boehringer Ingelheim, Ingelheim am Rhein, Germany

Jatin Arora	Boehringer Ingelheim, Ingelheim am Rhein, Germany

Ghadi Rai	Boehringer Ingelheim, Ingelheim am Rhein, Germany

Zhihao Ding	Boehringer Ingelheim, Ingelheim am Rhein, Germany

Lorenz Maier	Boehringer Ingelheim, Ingelheim am Rhein, Germany

Karsten Quast	Boehringer Ingelheim, Ingelheim am Rhein, Germany

Francisco Herruzo	Boehringer Ingelheim, Ingelheim am Rhein, Germany

Daniel Lopez	Boehringer Ingelheim, Ingelheim am Rhein, Germany

Marc Jung	Boehringer Ingelheim, Ingelheim am Rhein, Germany

Boris Bartholdy	Boehringer Ingelheim, Ingelheim am Rhein, Germany

Joseph Maranville	Celgene, Summit, NJ, United States/ Bristol Myers Squibb, New York, NY, United States

Shameek Biswas	Celgene, Summit, NJ, United States/ Bristol Myers Squibb, New York, NY, United States

Elmutaz Mohammed	Celgene, Summit, NJ, United States/ Bristol Myers Squibb, New York, NY, United States

Samir Wadhawan	Celgene, Summit, NJ, United States/ Bristol Myers Squibb, New York, NY, United States

Erika Kvikstad	Celgene, Summit, NJ, United States/ Bristol Myers Squibb, New York, NY, United States

Diana Chang	Genentech, San Francisco, CA, United States

Julie Hunkapiller	Genentech, San Francisco, CA, United States

Tushar Bhangale	Genentech, San Francisco, CA, United States

Natalie Bowers	Genentech, San Francisco, CA, United States

Rion Pendergrass	Genentech, San Francisco, CA, United States

Karen S King	GlaxoSmithKline, Brentford, United Kingdom

Padhraig Gormley	GlaxoSmithKline, Brentford, United Kingdom

Jimmy Liu	GlaxoSmithKline, Brentford, United Kingdom

Karsten Sieber	Janssen Research & Development, LLC, Spring House, PA, United States

Amy Hart	Janssen Research & Development, LLC, Spring House, PA, United States

Meijian Guan	Janssen Research & Development, LLC, Spring House, PA, United States

Shicheng Guo	Janssen Research & Development, LLC, Spring House, PA, United States

Matt Brauer	Maze Therapeutics, San Francisco, CA, United States

Jason Miller	Merck, Kenilworth, NJ, United States

Fabiana Farias	Merck, Kenilworth, NJ, United States

Jorge Del-Aguila	Merck, Kenilworth, NJ, United States

Kirill Shkura	Merck, Kenilworth, NJ, United States

Victor Neduva	Merck, Kenilworth, NJ, United States

Huilei Xu	Novartis, Basel, Switzerland

Amy Cole	Novartis, Basel, Switzerland

Jonathan Chung	Novartis, Basel, Switzerland

Jaison Jacob	Novartis, Basel, Switzerland

Katrina de Lange	Novartis, Basel, Switzerland

Jonas Zierer	Novartis, Basel, Switzerland

Xing Chen	Pfizer, New York, NY, United States

Åsa Hedman	Pfizer, New York, NY, United States

Franck Auge	Sanofi, Paris, France

Clement Chatelain	Sanofi, Paris, France

Deepak Rajpal	Sanofi, Paris, France

Dongyu Liu	Sanofi, Paris, France

Katherine Call	Sanofi, Paris, France

Tai-He Xia	Sanofi, Paris, France

Mitja Kurki	Institute for Molecular Medicine Finland, HiLIFE, University of Helsinki, Finland / Broad Institute, Cambridge, MA, United States

Samuli Ripatti	Institute for Molecular Medicine Finland, HiLIFE, University of Helsinki, Finland

Mark Daly	Institute for Molecular Medicine Finland, HiLIFE, University of Helsinki, Finland

Juha Karjalainen	Institute for Molecular Medicine Finland, HiLIFE, University of Helsinki, Finland

Aki Havulinna	Institute for Molecular Medicine Finland, HiLIFE, University of Helsinki, Finland

Juha Mehtonen	Institute for Molecular Medicine Finland, HiLIFE, University of Helsinki, Finland

Priit Palta	Institute for Molecular Medicine Finland, HiLIFE, University of Helsinki, Finland

Shabbeer Hassan	Institute for Molecular Medicine Finland, HiLIFE, University of Helsinki, Finland

Pietro Della Briotta Parolo	Institute for Molecular Medicine Finland, HiLIFE, University of Helsinki, Finland

Wei Zhou	Broad Institute, Cambridge, MA, United States

Mutaamba Maasha	Broad Institute, Cambridge, MA, United States

Shabbeer Hassan	Institute for Molecular Medicine Finland, HiLIFE, University of Helsinki, Finland

Susanna Lemmelä	Institute for Molecular Medicine Finland, HiLIFE, University of Helsinki, Finland

Manuel Rivas	University of Stanford, Stanford, CA, United States

Aarno Palotie	Institute for Molecular Medicine Finland, HiLIFE, University of Helsinki, Finland

Arto Lehisto	Institute for Molecular Medicine Finland, HiLIFE, University of Helsinki, Finland

Andrea Ganna	Institute for Molecular Medicine Finland, HiLIFE, University of Helsinki, Finland

Vincent Llorens	Institute for Molecular Medicine Finland, HiLIFE, University of Helsinki, Finland

Hannele Laivuori	Institute for Molecular Medicine Finland, HiLIFE, University of Helsinki, Finland

Taru Tukiainen	Institute for Molecular Medicine Finland, HiLIFE, University of Helsinki, Finland

Mary Pat Reeve	Institute for Molecular Medicine Finland, HiLIFE, University of Helsinki, Finland

Henrike Heyne	Institute for Molecular Medicine Finland, HiLIFE, University of Helsinki, Finland

Nina Mars	Institute for Molecular Medicine Finland, HiLIFE, University of Helsinki, Finland

Kimmo Palin	University of Helsinki, Helsinki, Finland

Javier Garcia-Tabuenca	University of Tampere, Tampere, Finland

Harri Siirtola	University of Tampere, Tampere, Finland

Tuomo Kiiskinen	Institute for Molecular Medicine Finland, HiLIFE, University of Helsinki, Finland

Jiwoo Lee	Institute for Molecular Medicine Finland, HiLIFE, University of Helsinki, Finland / Broad Institute, Cambridge, MA, United States

Kristin Tsuo	Institute for Molecular Medicine Finland, HiLIFE, University of Helsinki, Finland / Broad Institute, Cambridge, MA, United States

Amanda Elliott	Institute for Molecular Medicine Finland, HiLIFE, University of Helsinki, Finland / Broad Institute, Cambridge, MA, United States

Kati Kristiansson	THL Biobank / The National Institute of Health and Welfare Helsinki, Finland

Mikko Arvas	Finnish Red Cross Blood Service / Finnish Hematology Registry and Clinical Biobank, Helsinki, Finland

Kati Hyvärinen	Finnish Red Cross Blood Service, Helsinki, Finland

Jarmo Ritari	Finnish Red Cross Blood Service, Helsinki, Finland

Olli Carpén	Helsinki Biobank / Helsinki University and Hospital District of Helsinki and Uusimaa, Helsinki

Johannes Kettunen	Northern Finland Biobank Borealis / University of Oulu / Northern Ostrobothnia Hospital District, Oulu, Finland

Katri Pylkäs	University of Oulu, Oulu, Finland

Eeva Sliz	University of Oulu, Oulu, Finland

Minna Karjalainen	University of Oulu, Oulu, Finland

Tuomo Mantere	Northern Finland Biobank Borealis / University of Oulu / Northern Ostrobothnia Hospital District, Oulu, Finland

Eeva Kangasniemi	Finnish Clinical Biobank Tampere **/** University of Tampere / Pirkanmaa Hospital District, Tampere, Finland

Sami Heikkinen	University of Eastern Finland, Kuopio, Finland

Arto Mannermaa	Biobank of Eastern Finland / University of Eastern Finland / Northern Savo Hospital District, Kuopio, Finland

Eija Laakkonen	University of Jyväskylä, Jyväskylä, Finland

Dhanaprakash Jambulingam	University of Turku, Turku, Finland

Venkat Subramaniam Rathinakannan	University of Turku, Turku, Finland

Nina Pitkänen	Auria Biobank / University of Turku / Hospital District of Southwest Finland, Turku, Finland

**Biobank directors**

Lila Kallio	Auria Biobank / University of Turku / Hospital District of Southwest Finland, Turku, Finland

Sirpa Soini	THL Biobank / The National Institute of Health and Welfare Helsinki, Finland

Jukka Partanen	Finnish Red Cross Blood Service / Finnish Hematology Registry and Clinical Biobank, Helsinki, Finland

Eero Punkka	Helsinki Biobank / Helsinki University and Hospital District of Helsinki and Uusimaa, Helsinki

Raisa Serpi	Northern Finland Biobank Borealis / University of Oulu / Northern Ostrobothnia Hospital District, Oulu, Finland

Sanna Siltanen	Finnish Clinical Biobank Tampere **/** University of Tampere / Pirkanmaa Hospital District, Tampere, Finland

Veli-Matti Kosma	Biobank of Eastern Finland / University of Eastern Finland / Northern Savo Hospital District, Kuopio, Finland

Teijo Kuopio	Central Finland Biobank / University of Jyväskylä / Central Finland Health Care District, Jyväskylä, Finland

**FinnGen Teams**

**Administration**

Anu Jalanko	Institute for Molecular Medicine Finland, HiLIFE, University of Helsinki, Finland

Huei-Yi Shen	Institute for Molecular Medicine Finland, HiLIFE, University of Helsinki, Finland

Risto Kajanne	Institute for Molecular Medicine Finland, HiLIFE, University of Helsinki, Finland

Mervi Aavikko	Institute for Molecular Medicine Finland, HiLIFE, University of Helsinki, Finland

**Analysis**

Mitja Kurki	Institute for Molecular Medicine Finland, HiLIFE, University of Helsinki, Finland / Broad Institute, Cambridge, MA, United States

Juha Karjalainen	Institute for Molecular Medicine Finland, HiLIFE, University of Helsinki, Finland

Pietro Della Briotta Parolo	Institute for Molecular Medicine Finland, HiLIFE, University of Helsinki, Finland

Arto Lehisto	Institute for Molecular Medicine Finland, HiLIFE, University of Helsinki, Finland

Juha Mehtonen	Institute for Molecular Medicine Finland, HiLIFE, University of Helsinki, Finland

Wei Zhou	Broad Institute, Cambridge, MA, United States

Masahiro Kanai	Broad Institute, Cambridge, MA, United States

Mutaamba Maasha	Broad Institute, Cambridge, MA, United States

**Clinical Endpoint Development**

Hannele Laivuori	Institute for Molecular Medicine Finland, HiLIFE, University of Helsinki, Finland

Aki Havulinna	Institute for Molecular Medicine Finland, HiLIFE, University of Helsinki, Finland

Susanna Lemmelä	Institute for Molecular Medicine Finland, HiLIFE, University of Helsinki, Finland

Tuomo Kiiskinen	Institute for Molecular Medicine Finland, HiLIFE, University of Helsinki, Finland

L. Elisa Lahtela	Institute for Molecular Medicine Finland, HiLIFE, University of Helsinki, Finland

**Communication**

Mari Kaunisto	Institute for Molecular Medicine Finland, HiLIFE, University of Helsinki, Finland

**E-Science**

Elina Kilpeläinen	Institute for Molecular Medicine Finland, HiLIFE, University of Helsinki, Finland

Timo P. Sipilä	Institute for Molecular Medicine Finland, HiLIFE, University of Helsinki, Finland

Oluwaseun Alexander Dada	Institute for Molecular Medicine Finland, HiLIFE, University of Helsinki, Finland

Awaisa Ghazal	Institute for Molecular Medicine Finland, HiLIFE, University of Helsinki, Finland

Anastasia Shcherban	Institute for Molecular Medicine Finland, HiLIFE, University of Helsinki, Finland

Rigbe Weldatsadik	Institute for Molecular Medicine Finland, HiLIFE, University of Helsinki, Finland

**Genotyping**

Kati Donner	Institute for Molecular Medicine Finland, HiLIFE, University of Helsinki, Finland

Timo P. Sipilä	Institute for Molecular Medicine Finland, HiLIFE, University of Helsinki, Finland

**Sample Collection Coordination**

Anu Loukola	Helsinki Biobank / Helsinki University and Hospital District of Helsinki and Uusimaa, Helsinki

**Sample Logistics**

Päivi Laiho	THL Biobank / The National Institute of Health and Welfare Helsinki, Finland

Tuuli Sistonen	THL Biobank / The National Institute of Health and Welfare Helsinki, Finland

Essi Kaiharju	THL Biobank / The National Institute of Health and Welfare Helsinki, Finland

Markku Laukkanen	THL Biobank / The National Institute of Health and Welfare Helsinki, Finland

Elina Järvensivu	THL Biobank / The National Institute of Health and Welfare Helsinki, Finland

Sini Lähteenmäki	THL Biobank / The National Institute of Health and Welfare Helsinki, Finland

Lotta Männikkö	THL Biobank / The National Institute of Health and Welfare Helsinki, Finland

Regis Wong	THL Biobank / The National Institute of Health and Welfare Helsinki, Finland

**Registry Data Operations**

Minna Brunfeldt	THL Biobank / The National Institute of Health and Welfare Helsinki, Finland

Hannele Mattsson	THL Biobank / The National Institute of Health and Welfare Helsinki, Finland

Kati Kristiansson	THL Biobank / The National Institute of Health and Welfare Helsinki, Finland

Susanna Lemmelä	Institute for Molecular Medicine Finland, HiLIFE, University of Helsinki, Finland

Sami Koskelainen	THL Biobank / The National Institute of Health and Welfare Helsinki, Finland

Tero Hiekkalinna	THL Biobank / The National Institute of Health and Welfare Helsinki, Finland

Teemu Paajanen	THL Biobank / The National Institute of Health and Welfare Helsinki, Finland

**Sequencing Informatics**

Priit Palta	Institute for Molecular Medicine Finland, HiLIFE, University of Helsinki, Finland

Kalle Pärn	Institute for Molecular Medicine Finland, HiLIFE, University of Helsinki, Finland

Mart Kals	Institute for Molecular Medicine Finland, HiLIFE, University of Helsinki, Finland

Shuang Luo	Institute for Molecular Medicine Finland, HiLIFE, University of Helsinki, Finland

Vishal Sinha	Institute for Molecular Medicine Finland, HiLIFE, University of Helsinki, Finland

**Trajectory**

Tarja Laitinen	Pirkanmaa Hospital District, Tampere, Finland

Mary Pat Reeve	Institute for Molecular Medicine Finland, HiLIFE, University of Helsinki, Finland

Marianna Niemi	University of Tampere, Tampere, Finland

Harri Siirtola	University of Tampere, Tampere, Finland

Javier Gracia-Tabuenca	University of Tampere, Tampere, Finland

Mika Helminen	University of Tampere, Tampere, Finland

Tiina Luukkaala	University of Tampere, Tampere, Finland

Iida Vähätalo	University of Tampere, Tampere, Finland

**Data protection officer**

Jyrki Pitkänen	Institute for Molecular Medicine Finland, HiLIFE, University of Helsinki, Finland

**FINBB - Finnish biobank cooperative**

Marco Hautalahti

Johanna Mäkelä

Sarah Smith

Tom Southerington
