## Supplemental Figures for "SIRPα controls CD47-dependent platelet clearance in mice and humans"

### Supplementary figures

PheWAS associations with CD47-BBX locus SNP  rs167924.

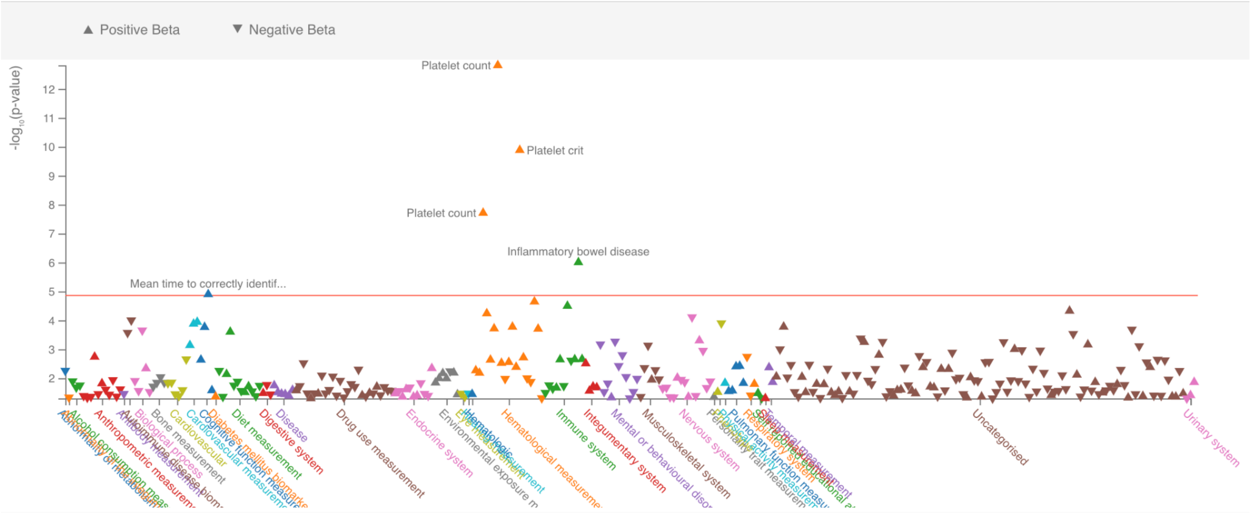


Platelet meta-analysis manhattan plot

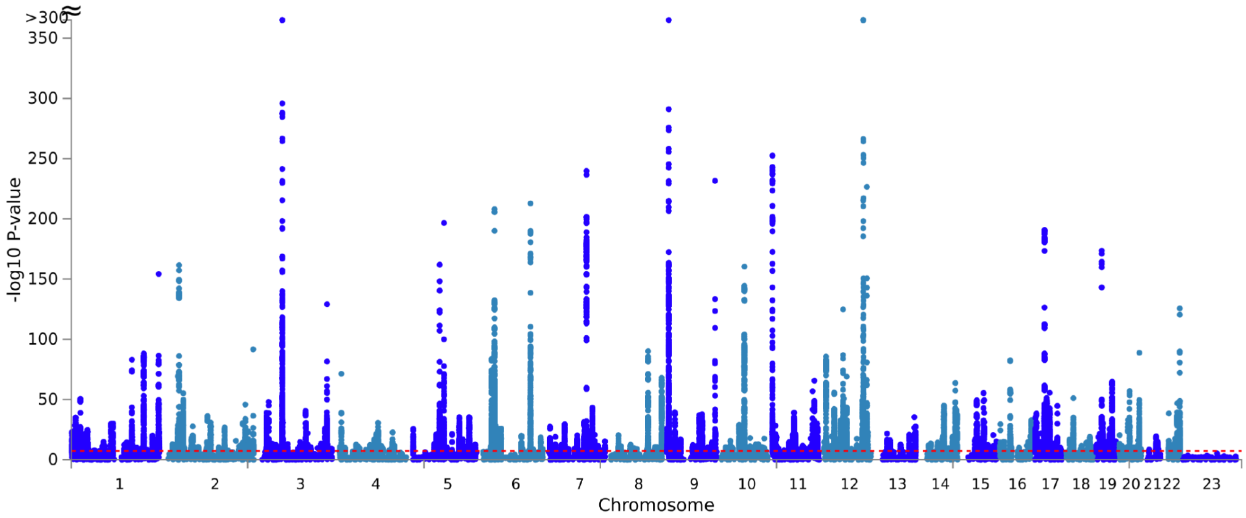
